# Changes in community composition determine recovery trajectories from multiple agricultural stressors in freshwater ecosystems

**DOI:** 10.1101/2020.11.03.366211

**Authors:** Francesco Polazzo, Talles Bruno Oliveira dos Anjos, Alba Arenas-Sánchez, Susana Romo, Marco Vighi, Andreu Rico

## Abstract

Pesticides have been identified worldwide as a threat for aquatic biodiversity due to their widespread use in agriculture and their capacity to reach freshwater ecosystems. Very little is known about the consequences of pesticide mixtures targeting different organism groups on community dynamics. Especially, how horizontal changes within one trophic level are propagated vertically across the food web has been rarely investigated. To get insight on the effects of pesticide mixtures on community dynamics, we performed a mesocosm experiment manipulating three common agricultural stressors: chlorpyrifos (an insecticide), diuron (an herbicide) and nutrients. The results of this study show that all stressors had significant effects on community composition, species richness and abundance. However, recovery trajectories and ecosystem functioning effects largely depended on the type of stressor as well as on post-disturbance trophic and non-trophic interactions. Effects of pesticides were generally recovered by the end of the experimental period when considering abundance, whereas community composition further departed from control systems. High nutrient loads led to a shift in community composition characterized by high taxa dominance and lower species richness, which in turn contributed to increased total organism abundance and reduced recovery times to pesticide exposure. We found interactions between the tested stressors to be significant only few times at the community level, while interactions were more common at the population level. Our findings indicate that management of freshwater ecosystems should consider pre-disturbance community composition and long-term changes in interactions across different organism groups to set effective protection measures.

## Introduction

Multiple anthropogenic stressors constantly impair ecosystems (Millennium Ecosystem Assessement 2005). Particularly, the freshwater realm has been highlighted as heavily impacted by a large number of disturbances, such as chemical pollution (Arenas-Sánchez et al., 2019; Rico et al., 2019), eutrophication (Woodward, 2012) and altered flow regimes (Matthaei, Piggott, & Townsend, 2010). The majority of studies assessing the effects of multiple stressors usually describe changes in community composition following disturbance (Barmentlo et al., 2018; Bray et al., 2019; Halstead et al., 2014; Matthaei et al., 2010; Piggott et al., 2015; Piggott, Townsend, & Matthaei, 2015), and only few focused on describing post-disturbance recovery trajectories (Arenas-Sánchez et al., 2019; Barmentlo et al., 2019; Rico et al., 2018). However, whether changes in community composition in one trophic level (or organism group) influence ecological responses of other trophic levels and their recovery dynamics is rarely studied.

Pesticides reaching freshwater ecosystems act as selective stressors targeting different organisms based on their physicochemical properties and toxicological mode of action, leading to non-random effects on communities (De Laender et al., 2016). Pesticides are known to reduce diversity (Bray et al., 2019), produce shifts in trophic interactions (Schrama et al., 2017) and affect ecosystem functioning (Spaak et al., 2017). Linking knowledge about the toxicological properties of pesticides and the implications for the whole food web is crucial to predict the net effects of complex chemical mixtures on aquatic ecosystems. The impact of pesticide mixtures on aquatic communities must be evaluated considering two different types of effects: the direct toxic effects, which may be estimated according to the models of Concentration Addition or Independent Action (Greco et al., 1992) for chemicals with the same toxicological mode of action or different, respectively; and the indirect ecological effects produced by compounds acting on different species groups, for example, an insecticide and an herbicide. To reach this objective, there is a need to unravel how different chemical stressors influence ecological dynamics horizontally within one trophic level (i.e., non-trophic interactions (Kéfi et al., 2015)) and vertically across different trophic levels (i.e, trophic interactions; Terry, Morris, & Bonsall, 2017). Moreover, we need to better understand how stressors affecting complex food webs drive changes in ecosystem functioning, and what the role of biodiversity is on preserving this. Most of the studies addressing biodiversity-ecosystem functioning relationships under experimental conditions focus on relatively simple food webs or isolated organism groups (Baert et al. 2016; Hooper et al. 2012; Spaak et al. 2017), excluding trophic and non-trophic interactions propagated vertically across the food web (but see Radchuk et al. 2016), and failing to provide effective ecosystem management actions at the ecosystem level. Nevertheless, these studies have highlighted that, under stress conditions, higher species richness does not always corresponds to higher levels of ecosystem functioning (Baert, Eisenhauer, Janssen, & De Laender, 2018; Spaak et al., 2017). Alternatively, it seems that community structure and the relative abundance of individuals in single populations determine the overall function of a system (Spaak et al., 2017; Van de Perre et al., 2018). Therefore, dominance of highly productive species could increase the degree of functioning carried out by a specific organism group, even though species richness remains the same or even decreases (Flöder & Hillebrand, 2012).

Community composition is actively modified by the underlying environmental conditions of ecosystems (Eigemann, Hilt, Salka, & Grossart, 2013; Lange, Liess, Piggott, Townsend, & Matthaei, 2011; Marshall, Karst, Nielsen, & Jørgensen, 2018; Robinson et al., 2018; Xun et al., 2015). Species composition and the relative abundance of species determine several ecological features such as resistance, recovery and resilience to disturbance (Griffiths et al., 2008; Vighi & Rico, 2018). Therefore, shifts in community composition imposed by changes in environmental conditions may also determine changes in single and multiple stressor effects. Nutrient enrichment is a common agricultural stressor, which contributes to eutrophication and to modify environmental conditions (Piggott, Townsend, and Matthaei 2015; Bray et al. 2019; Barmentlo et al. 2019). Increased nutrient concentrations can push ecosystems to a regime shift (Scheffer & Van Nes, 2007), modify community structure (Barmentlo et al. 2019) and decrease biodiversity, favouring a more homogenized community across a variety of ecosystems (Donohue et al., 2009; Isbell et al, 2013; Licursi, Gómez, & Sabater, 2016). Several studies have shown that eutrophication can reduce the impact of various stressors, from fine sediment addition (Matthaei et al., 2010; Piggott et al., 2015; Piggott, Townsend, & Matthaei, 2015) to chemicals (Halstead et al., 2014), by improving primary productivity, and generally increasing energy fluxes and biomass. Therefore, projections on the impacts of pesticides on biodiversity, ecosystem functioning and on post-disturbance recovery, should consider community composition scenarios that represent different trophic status of freshwater ecosystems.

Here, we address the importance of stressor type on post-disturbance trophic and non-trophic interactions by experimentally testing the single and combined effects of two widely used pesticides, with different toxicological properties, on community structure, resilience and recovery dynamics of macroinvertebrates, zooplankton and phytoplankton communities. To investigate the role of nutrients in these ecological dynamics, we tested stressors’ effects under two different pre-disturbance scenarios, one affected by nutrients enrichment and the other under natural trophic conditions using freshwater mesocosms. Moreover, we assessed the consequences of multiple stressor effects on trophic interactions, studying how changes in those interactions across organism groups determine different recovery trajectories. We also evaluated the implications of altered trophic relations for ecosystem functioning taking chlorophyll-a concentration as a proxy. The chemical stressors selected were the phenylurea herbicide diuron, which is known to inhibit the photosystem-II of primary producers, and the organophosphate insecticide chlorpyrifos, which inhibits the acetylcholinesterase enzymatic activity and is highly toxic to aquatic arthropods (i.e., crustaceans and insects).

Our main objective was to highlight the relevance of stressor type and stressor combinations in modifying ecological interactions between organism groups and to understand how these changes influence recovery trajectories. Thus, we hypothesized that:

H1: changes in community composition resulting from disturbance are longer lasting when they involve shifts in trophic or non-trophic interactions.
H2: change in populations’ density determined by stress and consequent shifts in community composition, rather than changes in species richness, determine alterations in ecosystem functioning. H3: continuous exposure to high nutrient concentrations push freshwater communities towards a regime shift, characterized by modified composition, reduced diversity and potentially leading to change in ecosystem functioning and resilience to chemical stress.

To our knowledge, this is one of the first attempts to study a complex stressors combination on a realistic species assemblage, investigating the response of three different trophic levels and using post-disturbance dynamics to provide mechanistic understanding of multiple stressor effects. Overall, we show how food web theory supported by empirical data, and coupled with knowledge of stressor’s toxicological properties, can improve our understanding of multiple stressor effects across multiple levels of biological organization.

## Materials and methods

### Experimental design

An outdoor experiment was performed at the mesocosm facilities of the IMDEA Water Institute (Alcalá de Henares, Madrid, Spain) between May and July of 2019. Each mesocosm consisted of a PVC tank (diameter: 120 cm; water depth: 75 cm) initially filled with approximately 40 cm of silty-sand sediments and 850 L of water from an artificial lagoon. Mesocosms were stocked with aquatic macrophytes (*Myriophyllum sp.* and *Elodea sp.*) and invertebrates collected from unpolluted water bodies in the vicinity of Alcalá de Henares. The biological community in the mesocosms was allowed to establish, and homogenized among experimental units, for two months prior to the start of the experiment.

A full factorial design (n=3), including chlorpyrifos (two levels: present and absent), diuron (two levels: present and absent) and nutrients (two levels: added, not added) was used in a randomized fashion, resulting in eight different treatments. Chlorpyrifos and diuron were applied at the concentration of 1 μg/L and 18 μg/L respectively. Those values are around 10 times the quality standards established by the Water Framework Directive for these priority substances (European Council, 2007), and represent peak exposure concentrations measured in edge of agricultural field surface waters (Thompson et al., 2016). Nutrients (P and N) were applied twice per week as a solution containing 1.820 g of NH4NO3 and 0.208 g of KH2PO4, which resulted in a nutrient addition of 750 μg/L of N and 75 μg/L of P, respectively. These nutrient levels correspond to an eutrophic/hypereutrophic system. Nutrient additions started 3 weeks before the pesticide application. Moreover, before the first nutrient addition, macrophytes were initially removed in order to create a phytoplankton dominated system, typically occurring in hypereutrophic shallow waters. The mesocosms that did not receive nutrients were considered oligo-mesotrophic based on nutrient analyses. In these systems, macrophytes were allowed to grow naturally.

### Chemicals application, sampling and analysis

Chlorpyrifos and diuron were applied as a single pulse on the 27^th^ of May 2019. Stock solutions of diuron and chlorpyrifos were prepared in 1 L of Milli-Q water and 50 mL of pure ethanol to improve the solubility. The solutions were poured over the mesocosm water surface. Additionally, 1 L of Milli-Q water with 50 mL of pure ethanol was added to the controls. Immediately after the application the mesocosms were stirred with a wooden stick to allow mixing and homogenization throughout the water column. The mesocosm water was sampled 2h after the pesticides’ application to evaluate possible differences between the nominal and the measured concentrations. Water samples were also taken on days 1, 3, 6, 10 after the chlorpyrifos application to assess the dissipation of the insecticide in the mesocosm water. Chlorpyrifos and diuron were measured in the chemical control mesocosms 2h and 10 days after the pesticides application and at the same sampling times, chlorpyrifos was measured in the mesocosms treated with diuron only. Diuron was measured in the mesocosms treated with it 2h after application and in days: 7, 21, 35 and 50 and at 2h and day 21 in the mesocosms treated with chlorpyrifos only. Depth-integrated water samples (1 L) were taken by means of a PVC tube. Water samples were introduced into amber glass bottles and immediately stored at −20 °C until analysis. The analysis of chlorpyrifos was performed using a gas chromatograph (GC) system (Agilent 7890A) coupled to a mass spectrometer (MS) with a triple quadrupole analyser (Agilent 7000 GC/MS Triple Quad). Diuron was analysed using a high-performance liquid chromatography system (Agilent Technologies 1200) coupled with a time-of-flight mass spectrometry (TOF-MS) (Agilent Technologies 6230). The dissipation coefficients (*k*) of chlorpyrifos and diuron were calculated by means of linear regression of the ln-transformed concentrations with the software Microsoft Excel version 2010 assuming first-order kinetics. The half-lives (DT50s) of the evaluated insecticides were calculated by Ln (2) divided by *k*.

### Water quality parameters

Water samples (0.5 L) were collected on days −5, 15, 30 and 50 relative to the application of the pesticides in order to analyse the concentrations of ammonia, nitrate, ortho-phosphate and total phosphorous. The total inorganic nitrogen concentration in the mesocosm water was calculated as the sum of the concentrations of ammonia and nitrate, assuming nitrite as negligible. Analysis of nutrients concentrations were performed by colorimetry following slightly modified methods described in APHA (2005). Temperature, dissolved oxygen, pH, conductivity and turbidity were measured by a multimeter (HANNA HI0194) in the morning (8 a.m.) and afternoon (7 p.m.) on the days −5, 7, 15, 30 and 50 relative to the application of the pesticides. Additional water samples (0.5 L) were taken on days −5, 15, 30 and 50 relative to the pesticides application to assess the concentration of chlorophyll-a. Chloropyll-a was measured following the protocol described in APHA (2005) and was used as a proxy of the primary productivity of the suspended microalgae community.

### Phytoplankton

Phytoplankton were sampled on days −5, 7, 15, 50 relative to the pesticides’ application. Depth-integrated water samples were taken with a PVC tube (six sub-samples per mesocosm mixed in a bucket). Next, 250 mL of this sample was introduced into glass amber bottles and 10% Lugol was added for preservation. Counting and identification were done to the lowest taxonomic resolution possible with an inverted microscope (Motic AE31) coupled to a Motic Plan x 100/1.25 Oil ∞/0.17 objective.

### Zooplankton

Zooplankton samples were taken on days −5, 7, 14, 30 and 50 relative to the application of pesticides. Six depth-integrated samples of approximately 1 L were collected from each mesocosm using a PVC tube and mixed together. Subsequently 5 L of the composed water sample were passed through a zooplankton net (55 μm) and concentrated to an approximate volume of 100 mL. The concentrated samples were fixed with Lugol’s iodine solution and stored in dark conditions at room temperature until identification and counting. Cladocera, Ostracoda and Copepoda were identified and counted in the entire zooplankton sample using an Olympus SZx2-TR30 stereomicroscope with a magnification of 20×. As for Rotifera and Copepoda nauplii, a subsample of 1 mL was taken and analysed using a binocular Olympus UCTR30-2 microscope with a magnification of 100×. Cladocera and Rotifera were identified up to the genus or species level. For Cyclopoida, a distinction was made between the adult stages (as the sum of true adults and copepodite stages) and the nauplius stages during counting and for further statistical analyses. Ostracoda appeared in relatively low abundances and were not further identified.

### Macroinvertebrates

Macroinvertebrates were sampled on days −32, 15, 30 and 50 relative to the application of the pesticides. In order to collect pelagic and benthic individuals, three sampling methods were used. First, a net (mesh size: 0.5 mm) was passed twice through the side of the mesocosms (in both directions) to catch the animals that were swimming or resting on the mesocosm’s wall. Second, two pebble baskets positioned over the sediment surface were collected, and third, two traps filled with macrophytes (*Elodea sp.), Populus sp.* leaves and stones were collected from the sediment’s surface using a net. The invertebrates sampled from each mesocosm with the three sampling methods were pooled together, identified and counted alive. Afterwards, the invertebrates were returned to their original mesocosms together with the colonizing pebble baskets and traps. The macroinvertebrate taxonomic identification was performed to the lowest practical resolution level making use of freshwater biology guides (e.g. Tachet et al., 2000).

### Statistical analysis

To examine the effects of diuron, chlorpyrifos, nutrients and their possible interaction on the communities (phytoplankton, zooplankton, macroinvertebrates), we used permutational multivariate analysis of variance (PERMANOVA) with 999 permutations (Anderson, 2001). Homogeneity of data dispersion across the different treatments was tested using a distance-based dispersion test, followed by a permutation-based test of multivariate homogeneity of group dispersions (variances). Single-taxon contribution to the overall Bray-Curtis dissimilarity and cumulative contribution were obtained by performing Similarity Percentages (SIMPER) to identify the taxa with the highest contribution to the variation in the assemblage composition observed in the PERMANOVA. To further explore the effects of treatments on the experimental communities, we analysed single taxon abundance response for some relevant taxa, indicated by the SIMPER analysis (the three taxa explaining most of the variation for each treatment), using a three-way ANOVA. We tested normal distribution and homogeneity of variances using the Shapiro-Wilk test combined with Quantile-Quantile plots and Levene’s test, respectively. When the assumptions were not met, the data were either log or square root transformed. Three-way ANOVA was used to study the effects of the treatments on water quality parameters as well. Furthermore, we applied some diversity indexes to investigate the effects of single and combined stressors on communities. We calculated species richness, Shannon’s index and Berger-Parker dominance index using the R packages “BiodiversiytR” and “lawstat”. Differences among indexes’ values caused by treatments were analysed by means of a three-way ANOVA. We considered statistically significant differences when the calculated p-value was <0.05. All statistical analyses and figures were created using R (version 3.5.1) in RStudio Team (2018).

## Results

### Pesticides concentrations and physiochemical parameters

Two hours after application, measured concentrations of chlorpyrifos and diuron were very close to the nominal concentrations, indicating an adequate application of the test compounds (see Supporting Information, Table S1 and S2). Mean calculated DT50s for chlorpyrifos and diuron were 2.4 days and 39 days, respectively (Table S1 and S2). Overall, we observed a faster pesticides’ dissipation in the mesocosms treated with nutrients (up to about 30% for chlorpyrifos and 20% for diuron).

Physicochemical parameters responded consistently to the treatments (Table S3 and S4). Nutrients were significantly higher in mesocosms undergoing nutrients addition, and in those systems, chlorophyll-a, oxygen concentration and pH were also significantly higher. Diuron application, significantly decreased all parameters related to photosynthetic activity: oxygen concentration and pH, with the exception of chlorophyll-a concentration (Figure 1, Tables S3 and S4).

**Figure 1.**
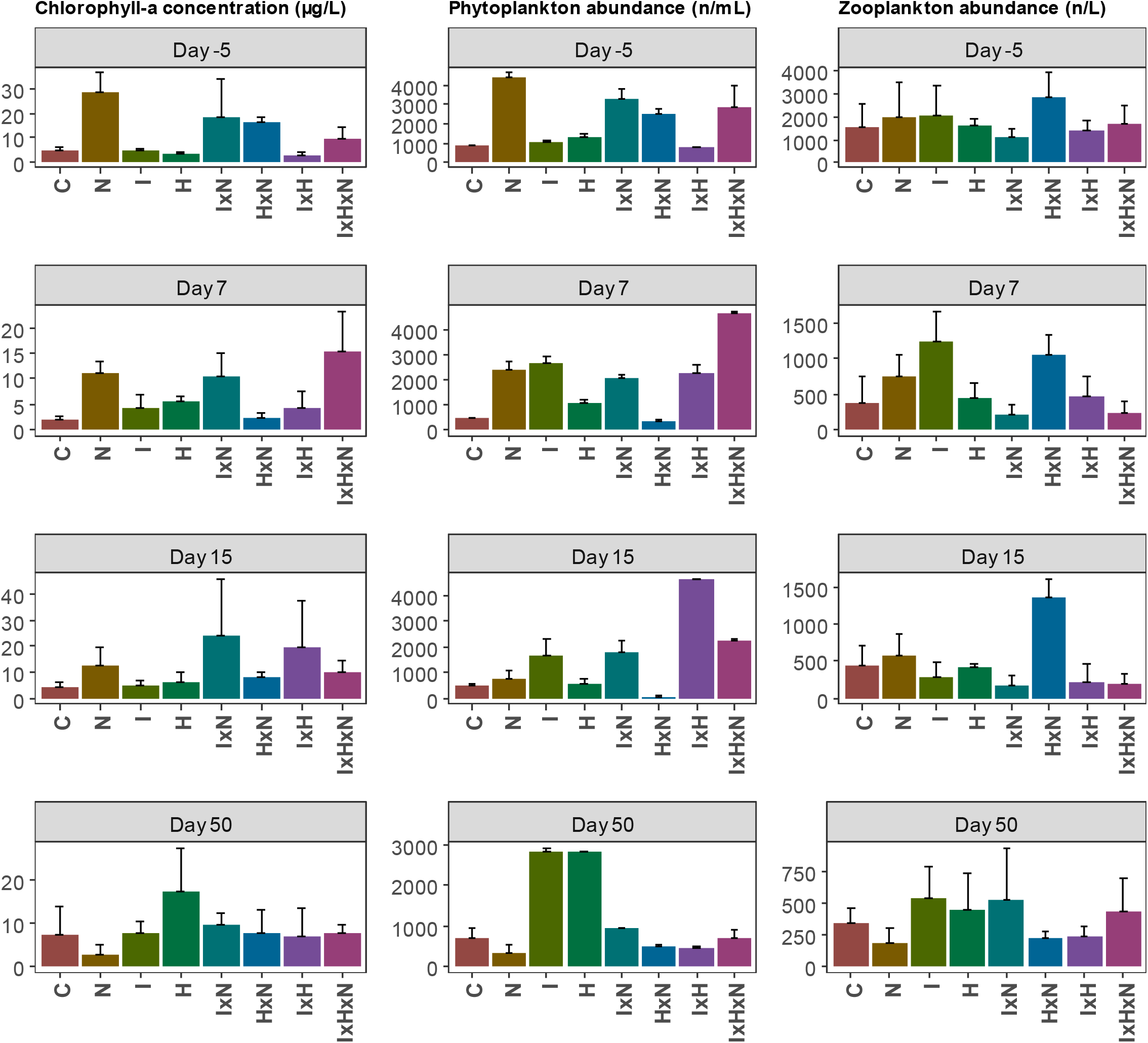
Chlorophyll-a concentration (μg/L), total abundance of phytoplankton (individual/mL) and total abundance of zooplankton (individual/L) across the experimental time. Abundance and concentrations are expressed as treatment average for each sampling day with their standard deviation. Zooplankton abundance on day 30 is provided in SI figure S3: controls; N: nutrients; I: insecticide/chlorpyrifos; H: herbicide/diuron.

### Phytoplankton responses

Sixty-eight phytoplankton taxa were monitored in the mesocosms during the experimental period. The phytoplankton community was formed by Chlorophyta (24 taxa), Desmids (6 taxa), Dinophyta (1 taxon), Cryptophyta (3 taxa), Diatoms (17 taxa), Cyanobacteria (13 taxa) and Euglenophyta (4 taxa). The PERMANOVA analysis shows that the application of nutrients significantly modified the phytoplankton community composition during the whole experimental period (Table 1). Mesocosms treated with nutrients were generally characterized by high total abundance, and lower species richness towards the end of the experiment (Figure 1, Table S5). Chlorpyrifos significantly affected community composition, starting from day 15 onwards (Table 1). In the mesocosms treated with chlorpyrifos, abundance was significantly increased, particularly Chlorophyta (the most abundant taxonomic group) which significantly increased on days 7 and 50 (Figure S1, S2, Table S5 and S6). Diuron modified community composition in every sampling day after its application (Table 1). Total abundance was never negatively impacted, but species richness decreased (Table S5). Diatoms, Cryptophyta and Cyanobacteria were the most affected taxonomic groups (Table S6). On day 50, the interaction between diuron and nutrients was significant, which indicates an effect on community composition characterized by a significant decrease of Diatoms (Figure S1, Table S6). Other stressor interactions were not significant.

**Table 1.** Summary of p-values resulting from the Permutational Analysis of Variance (PERMANOVA) based on Bray - Curtis dissimilarities using abundance for phytoplankton, zooplankton and macroinvertebrates communities for the period pre (Day −5 and −32) and post-pesticide application (Days 7, 15, 30 and 50). Significant (p < 0.05) of the treatment are shown in bold. ‘***’ p<0.001, ‘**’ p<0.01, ‘*’p< 0.05. N: nutrients; I: insecticide/chlorpyrifos; H: herbicide/diuron.

|  | N | I | H | IxN | HxN | IxH | IxHxN |
| --- | --- | --- | --- | --- | --- | --- | --- |
| **Phytoplankton** |  |  |  |  |  |  |  |
| D-5 | **0.023 \*** | 0.517 | 0.904 | 0.480 | 0.431 | 0.360 | 0.680 |
| D7 | **0.048 \*** | 0.213 | **<0.001 \*\*\*** | 0.650 | 0.783 | 0.112 | 0.501 |
| D15 | **0.007 \*\*** | **<0.001 \*\*\*** | **<0.001 \*\*\*** | 0.421 | 0.199 | 0.712 | 0.461 |
| D50 | **0.003 \*\*** | **0.007 \*\*** | **0.006 \*\*** | 0.566 | **0.002 \*\*** | 0.566 | 0.293 |
| **Zooplankton** |  |  |  |  |  |  |  |
| D-5 | 0.178 | 0.941 | 0.483 | 0.632 | 0.162 | 0.578 | 0.647 |
| D7 | **0.011 \*** | **0.001 \*\*\*** | 0.649 | **0.037 \*** | 0.169 | **0.037 \*** | 0.183 |
| D15 | **<0.001 \*\*\*** | **0.001 \*\*\*** | 0.232 | 0.129 | 0.050 | 0.846 | 0.516 |
| D30 | **0.025 \*** | **0.001 \*\*\*** | **0.011 \*** | 0.975 | 0.222 | 0.284 | 0.981 |
| D50 | **0.014 \*** | **0.002 \*\*** | 0.265 | 0.514 | 0.514 | **0.044 \*** | 0.265 |
| **Macroinvertebrates** |  |  |  |  |  |  |  |
| D-32 | 0.883 | 0.3 | 0.442 | 0.01 | 0.952 | 0.57 | 0.273 |
| D15 | **0.041 \*** | **<0.001 \*\*\*** | **0.003 \*\*** | 0.349 | 0.071 | 0.079 | 0.407 |
| D30 | 0.121 | 0.054 | **<0.001 \*\*\*** | 0.405 | 0.642 | 0.263 | 0.076 |
| D50 | **0.005 \*\*** | 0.628 | 0.203 | 0.709 | 0.865 | 0.071 | 0.087 |

### Zooplankton responses

Twenty-two zooplankton taxa were monitored during the experimental period. The community was composed of Cladocera (5 taxa), Copepoda (2 taxa), Rotifera (14 taxa) and Ostracoda (1 taxon). Nutrients addition caused significant changes in the zooplankton community structure on every sampling day (Table 1). High nutrient concentrations resulted in some cases in increased abundances (Figure 1, Table S7), but also in a reduction of species richness and increased values of species dominance (Table S7). Particularly, some Rotifera species (*Testudinella sp* and *Polyarthra sp*) were severely impaired by the nutrients treatment (Table S8). Chlorpyrifos also shifted community composition in each sampling day after its application (Table 1). In the mesocosms treated with chlorpyrifos, zooplankton abundance was low until day 15, as were species richness and the Shannon diversity index, while taxa dominance was higher (Table S7). From day 30 to the end of the experiment, abundance levels were equal or higher compared to the controls, but richness, diversity and dominance were still dissimilar to the control mesocosms (Table S7). Chlorpyrifos caused a drastic reduction in Cladocera abundance (Figure S4 and S5), while the more tolerant Copepoda increased their abundance.

Significant interactions between stressors were detected for the zooplankton community composition (Table 1). The mixture of pesticides determined significant changes in community composition, associated to a reduced richness and lower abundance values. In the mesocosms exposed to the herbicide and insecticide, total abundance was negatively affected (Figure 1). Copepoda were more impacted in the first half of the experiment (day 7 and 14, Figure S5) and they recovered by the end of the experimental period. On the other hand, Cladocera abundance was low throughout the whole experiment in the mixture treatment (Figure S5).

### Macroinvertebrate responses

Thirty-eight macroinvertebrate taxa were monitored in the mesocosms during the experimental period. The macroinvertebrate community was formed by insects (26 taxa), molluscs (5 taxa), annelids (3 taxa), platyhelminths (2 taxa), a crustacean (1 taxon) and an arachnid (1 taxon). Nutrients addition caused a shift in macroinvertebrates community structure on day 15 and 50 (Table 1). The eutrophic conditions were generally associated with higher total abundance (Figure 2). However, by the end of the experimental period, high nutrient concentrations led to a decrease in species richness and to a community dominated by fewer species (Table S9). Chlorpyrifos had strong significant effects on the macroinvertebrates’ community structure in the first half of the experiment (Table 1). The community impaired by the insecticide showed reduced total abundance (Figure 2) and species richness (Table S9). Some sensitive taxa were temporally extinct (Ephemeroptera, Figure S7) and generally most of the insect taxa were significantly reduced (Table S10). However, more tolerant taxa with a simple nervous system (Annelida), increased their abundance (Table S10, Figure S6). On day 30, chlorpyrifos did not have any more a significant effect on community structure (Table 1), nor any deviation from the controls was noticed regarding total abundance or diversity indices (Table S9). The macroinvertebrate community treated with chlorpyrifos no longer deviated from controls at the end of the experiment for any compositional or diversity matrix.

**Figure 2.**
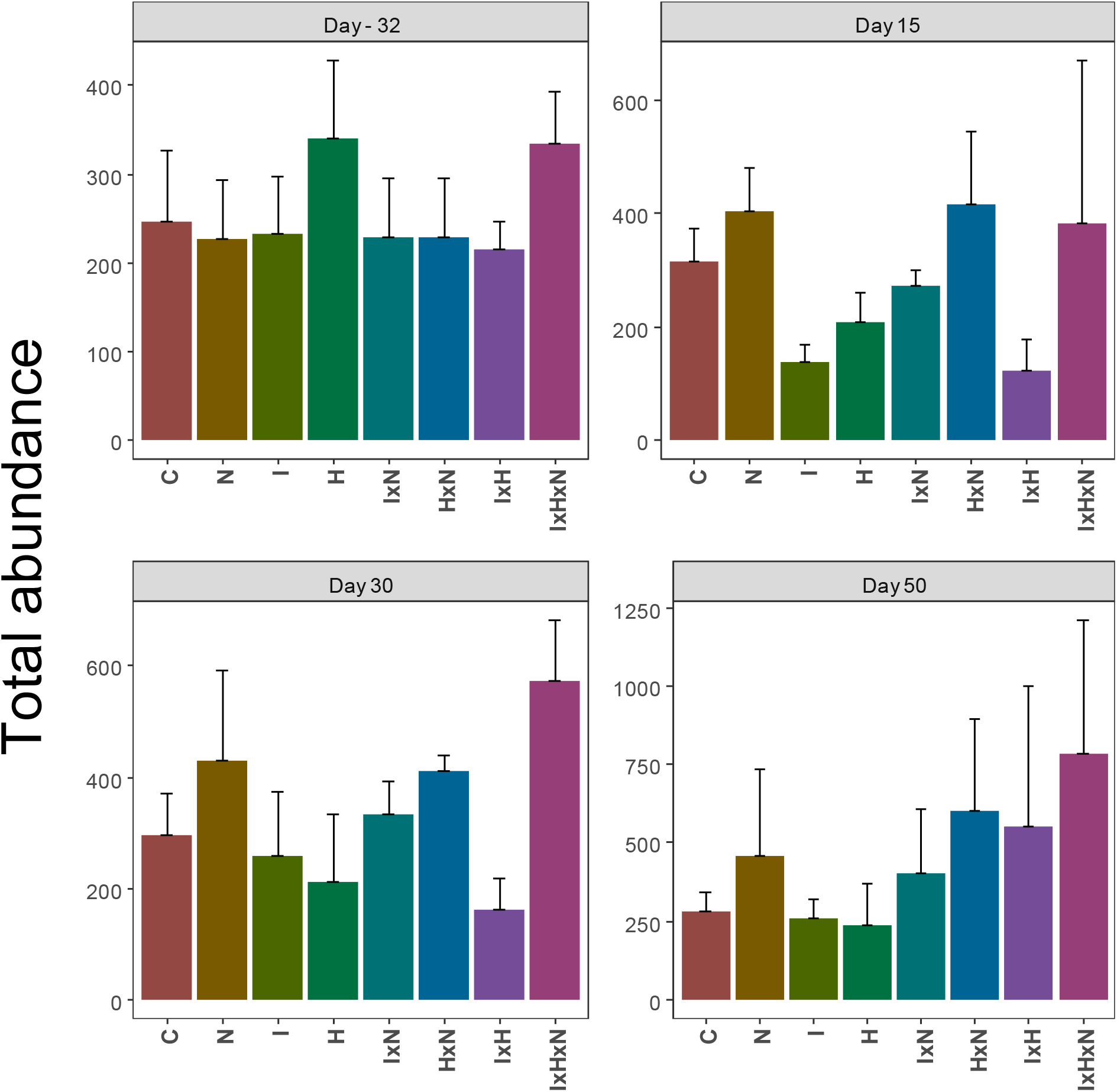
Total abundance of macroinvertebrates across the experimental period. Abundance is expressed as treatment average for each sampling day with their standard deviation. C: controls; N: nutrients; I: insecticide/chlorpyrifos; H: herbicide/diuron.

**Figure 3.**
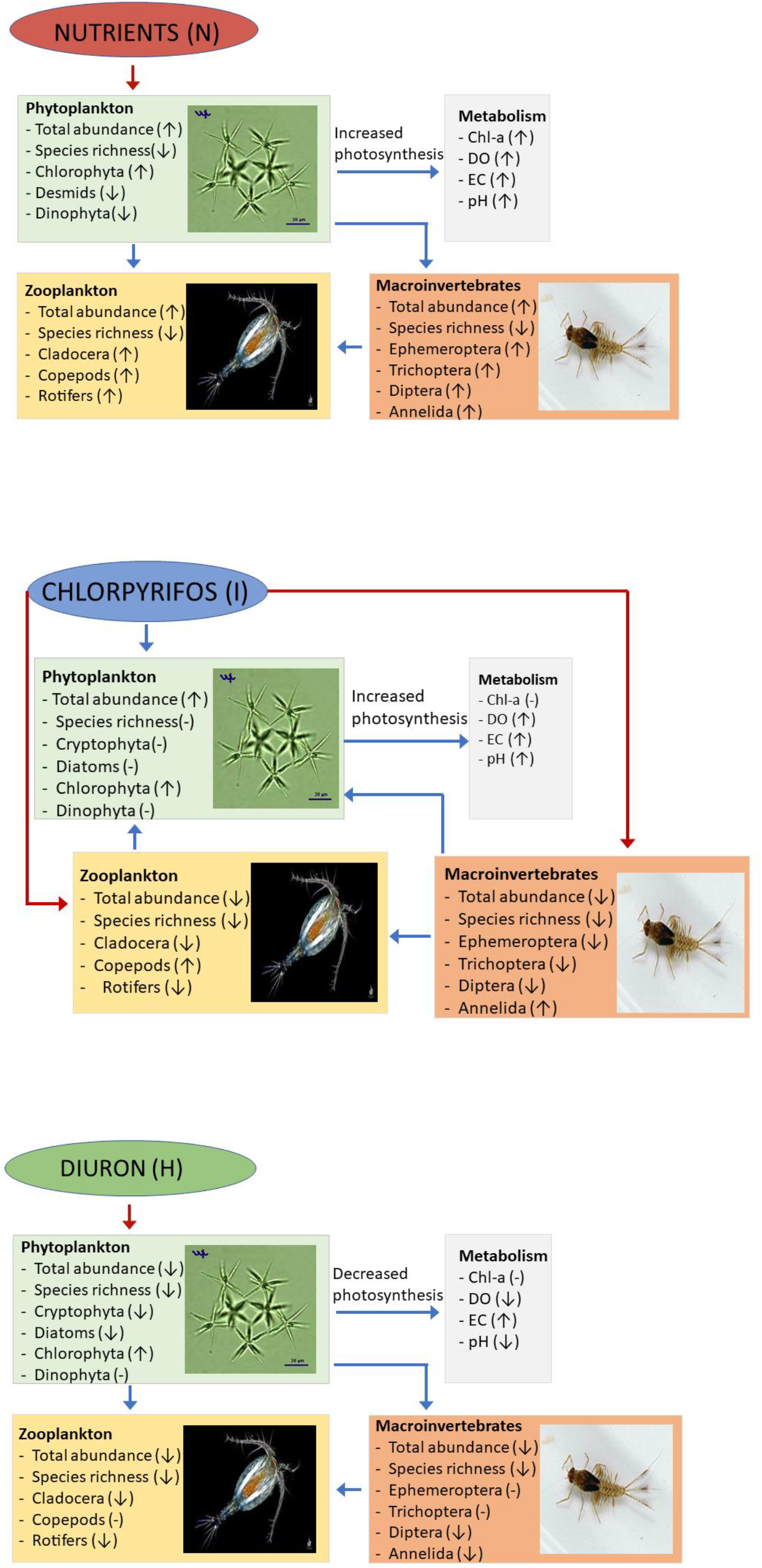
Conceptual map of the single effects of nutrients, chlorpyrifos and diuron. Arrows indicate a stressor-related increase (↟) or decrease (↓), (-) indicates no effect. Red lines connecting the boxes represent direct effects, and blue lines represent indirect effects. Conceptual maps showing the effects of multiple stressors are shown in Fig. S7.

Interactions between stressors were not significant at the community level, but there were significant stressors’ interactions at the population level. The Diptera taxa Chironomini and Orthocladiinae were negatively impaired by the joint effect of diuron and chlorpyrifos, whereas *Dugesia* sp. increased in abundance almost every sampling day (Table S10).

## Discussion

In this work we show how different pesticides affect sensitive communities in terms of species composition, richness, dominance and abundance depending on their toxicological properties. In H1, we hypothesized that changes in community composition resulting from disturbance are generally longer lasting when shifts in trophic and non-trophic interactions are involved. Disturbance often results in changes of one taxon abundance which can lead to changes in another taxon abundance (Tylianakis, Didham, Bascompte, & Wardle, 2008). Studies show that this mechanism holds horizontally as well as vertically, influencing several ecological processes (Gilbert et al., 2014; Zhao et al., 2019). Despite the pulsed stress characteristics of chlorpyrifos, it contributed to modify zooplankton’s community structure during the whole experimental period. At the end of the experiment, zooplankton community composition further deviated from the control systems. The missed compositional recovery of zooplankton reflects two main ecological stress-related features of the community: (a) different organism sensitivities to the insecticide and (b) changes in the strength of non-trophic interactions (competition). In our experiment, the insecticide disturbance caused the decline of the most sensitive taxa (Cladocera), allowing the more tolerant taxa (Copepoda) to thrive, shifting the abundance ratio of competing species. The post-disturbance disbalance in relative densities of competing species determined an alternative dominance state (*sensu* Fukami & Nakajima, 2011). Copepoda became the dominant species through competitive exclusion and niche pre-emption after Cladocera’s decline (Fukami, 2015). Supporting this argument, systems treated with chlorpyrifos (as a single stressor or as a mixture with diuron) showed the lowest abundance of Cladocera across all treatments in every sampling day (Figure S4), even when the insecticide concentration was negligible (day 50) and not expected to cause any harm to aquatic invertebrates (Huang et al., 2020). The establishment of a new dominant taxon due to a shift in non-trophic interactions strength led to a long-lasting modification of community structure (H1).

Chlorpyrifos was not expected to have a direct effect on primary producers at the tested concentration (Huang et al., 2020). Yet, phytoplankton community composition resulted altered during the whole experimental period in the insecticide treatment. We argue that the monitored changes in phytoplankton composition were driven by the above explained changes in grazers relative abundances and composition. Different zooplankton taxa have been shown to have different selective grazing behaviors (Fulton & Paerl, 1988; Hambright, Hairston, Schaffner, & Howarth, 2007) which can shape primary producers composition (Hillebrand et al., 2007; Müren, Berglund, Samuelsson, & Andersson, 2005; O’Connor, Piehler, Leech, Anton, & Bruno, 2009). In our systems, the reduction in cladoceran abundance and the inverse response of Copepoda, changed the selective grazing pressure exerted by the two taxa on phytoplankton. The shift in consumers’ community structure changed the nature of the top-down regulation (trophic interaction) which, in turn, actively shaped the community structure of primary producers. Thus, our results show that, even though a disturbance has a very specific target (i.e., pesticides with specific toxicological properties), indirect effects may propagate horizontally and vertically through the food web, determining different recovery trajectories in taxonomic groups not directly impaired by the stressor. The propagation of indirect effects through the food web could lead to deep alterations in the functioning of ecosystems, withdrawing consequences that are often overlooked, especially in the conventional ecological risk assessment of chemicals.

The herbicide-insecticide mixture caused a significant shift in zooplankton community composition, showing an interactive effect. The interaction between diuron and chlorpyrifos affected zooplankton through direct and indirect effects (Figure 5 and S7). The insecticide directly impaired some zooplanktonic crustaceans, causing the above-mentioned changes in community composition. Diuron shifted primary producers community structure and may have decreased the abundance of palatable phytoplankton species in the pesticides mixture treatment, thus reducing energy sources for grazers. The reduction in food resources contributed to increase the inter- and intra-specific competition in the zooplankton community. In support of this finding, zooplankton total abundance was decreased by almost 70% in the pesticides mixture treatment compared to the control on day 30, while in the insecticide only treatment, the total abundance was comparable to the control levels in the same day (118% of control). At the end of the experiment, the total abundance was still reduced by 30% in the pesticides mixture. Furthermore, the abundance of the less sensitive taxon Copepoda, was lower in the pesticides mixture treatment compared to the control and chlorpyrifos only treatment. The low abundance of the more tolerant organism group supports the hypothesis of food-limitation and consequent increase in intraspecific competition, which limit the recovery potential. Mechanistically, such results can be explained by the fact that long-lasting perturbation exerted by diuron (longer persistence) on primary producers’ composition did not allow zooplankton to recover. These results highlight that stressor interactions determine ecological dynamics of recovery (enhancing or delaying it) by modifying trophic and non-trophic interactions (H1), and should therefore be considered in ecological risk assessment when dealing with predictions of multiple stressor combinations.

In H2, we hypothesized that changes in community structure and single populations’ densities following disturbance determine fluctuations in ecosystem functioning, rather than the reduction or increase of diversity. Our results confirm H2 in multiple ways. The reduction in species richness (lower diversity) caused by nutrients addition did not reduce any functional parameter. In fact, oxygen concentration, chlorophyll-a and pH were increased in the systems treated with nutrients. The underpinning mechanism explaining the increase in phytoplankton function is a shift towards more tolerant species and an increase in functional replacement (and hence in the dominance effect). Increased abundance of species with higher per-capita contribution to function (Chlorophyta, particularly colony forming *Oocystis* sp.) sustained by high nutrients availability contributed to increase the overall system functioning. Despite the generally accepted positive correlation between diversity and ecosystem functioning, it is not the first time that deviations from this relationship are reported for phytoplankton (Baert et al., 2016; Spaak et al., 2017). Yet, our experiment is one of the first ones to provide evidence supporting this concept in a complex ecological food web, where not only horizontal diversity was altered, but also the total diversity of the system (different trophic levels) was reduced.

Phytoplankton was directly impaired by the herbicide diuron, which modified community composition and decreased richness. However, reduced richness and shift in composition did not result in a chlorophyll-a decrease when total abundance was not reduced. Indeed, chlorophyll-a concentration was not reduced in the systems treated with the herbicide. This result suggests that, if the initial number of species is high enough to assure functional redundancy, function (i.e. chlorophyll-a concentration) can be maintained invariant even after stress application and consequent reduction in diversity (Baert et al., 2016), as postulated by the insurance hypotheses (Yachi & Loreau, 1999). In addition, our results suggest that the shift in community composition was driven by a reduction in sensitive species (particularly small Diatoms), which in turn allowed more tolerant species to thrive (Van de Perre et al., 2018).

In the insecticide treatment, the total abundance of phytoplankton was increased, as primary producers benefited by the reduction in grazing pressure. Nevertheless, chlorophyll-a concentrations, although increased, were not significantly higher as compared to the control systems, showing discrepancy with the increase in total abundance. This can be attributed to the stronger interspecific competition for resources (light and nutrients mainly) caused by high abundance and to the selective grazing pressure carried out by Copepoda. This taxon has been shown to selectively graze on cells with higher chlorophyll-a content (Hambright et al., 2007). Therefore, selective grazing on species with high per-capita contribution to function, and increased competition may have contributed to limit chlorophyll-a concentration. The changes in phytoplankton composition were underpinned by an increased in species with low productivity, which explains the discrepancy between the high total abundance and the relatively low chlorophyll-a concentration. Thus, in an ecologically realistic species assemblage, the function performed by a group of organisms cannot be assessed without evaluating the effects of the biological interaction occurring between the organism group under investigation and the other trophic levels, as species interactions can strongly influence ecological functions.

In H3, we hypothesized that high nutrient contamination may alter community composition and modify their functioning and resilience to chemical stress. Here we demonstrate that nutrients’ addition modified the structure and diversity of all three communities examined in this study for the whole duration of it (also confirming H1). In eutrophic systems, the biotic response was rather consistent across different organism groups. Despite a general increase in total abundance, nutrients caused a decrease in species richness and an increase in species dominance. The phytoplankton increase in total abundance under eutrophic scenario persisted only up to day −5. Later, the initial increase in primary producers abundance was followed by a growth of the grazer zooplankton populations, which limited phytoplankton proliferation. High nutrient concentrations have been shown to alter the heterotrophs/autotrophs ratio in the long term, by increasing consumer biomass relative to producer biomass, and reducing the total biomass of the food web despite increases in primary productivity (O’Connor et al., 2009). The increase in primary productivity was reflected in our study by the water’s physiochemical parameters (pH and oxygen saturation), which were significantly higher in the mesocosms with nutrients’ addition.

Eutrophication is considered as one of the most important press disturbances to freshwater ecosystems (Woodward, 2012), particularly under Mediterranean conditions, which are subject to increasing water scarcity problems (Beklioğlu et al., 2020). Our study confirms that the continued exposure to high nutrients levels deeply modifies communities’ features, such as diversity and structure (Donohue et al., 2009; Hillebrand, Bennett, & Cadotte, 2008). Such changes have been proven to require several decades to recover after cessation of nutrients enrichment (Isbell et al., 2013). Moreover, similarly to Hillebrand et al. (2007), our results show that eutrophication increases species dominance. Dominance (or its complementary term, evenness) measures the traits-distribution of a community. High dominance of few species is associated with high density of a limited number of traits, constraining the overall functions an ecosystem performs. Yet, the traits distribution affects both the strength and sign of intra- and interspecific interactions, thus mediating the effects of these links on species coexistence (Hillebrand et al., 2008). Nutrients enrichment is often considered as an antagonistic factor against detrimental effects of other disturbance types in multiple stressor research (Barmentlo et al., 2019; Halstead et al., 2014; Matthaei et al., 2010; Piggott et al., 2015). Consistently, in our experiment, nutrients often reduced negative effects of pesticides, especially at the population level. Furthermore, high nutrients concentrations reduced bioavailability and accelerated the pesticides’ dissipation in the water column, reducing the overall toxic stress. Nevertheless, as described by several authors, eutrophication represents a primary threat to ecosystem functioning, community composition and biodiversity across ecosystems (Donohue et al., 2009; Isbell et al., 2013; Wåhlström et al., 2020; Woodward, 2012). Thus, the effects of high nutrients concentrations need to be careful evaluated as single disturbances as well as in interactions with other stressors to avoid misleading conclusions.

## Conclusions

This study shows that pesticides affecting different aquatic organism groups may result in different direct and indirect effects, which drive changes in community composition that differ among ecosystems with varying nutrient availability and eutrophic conditions. Such modified community compositions drive changes in trophic and non-trophic interaction, which, in turn, determine dissimilar recovery patterns to pesticides. Generally, we found that high nutrient availability reduces recovery times of phytoplankton, zooplankton and macroinvertebrates to the single and combined exposure to pesticides with different toxicological properties by increasing primary productivity and biomass. The pesticide dissipation was found to be faster under eutrophic conditions, which may also contribute to higher opportunities for recovery. However, we have shown that eutrophication contributes to homogenize freshwater communities’ composition and results in reduced biodiversity. This study also reveals that total organism abundance is generally recovered faster than community composition, regardless of stressor type. We also highlight that compositional recovery can be impaired in the long term, when trophic interactions are modified by perturbations. Post-disturbance trophic interactions can lead to competitive exclusion and can push communities to an alternative state. Furthermore, the present study provides experimental evidence that reduction in diversity is not always followed by a decline in ecosystem functioning. Moreover, we show that different stressors can have very little effects on ecosystem functioning in species rich systems.

Overall, the experimental data strongly support our initial hypotheses. Mechanistic understanding of stressors and trophic links on ecosystem functioning proposed in this study may have implications for ecosystem management and ecological risk assessment in the future. Indeed, we propose that pesticides’ effects on trophic relations across the food web should be considered to predict multiple stressor effects, since it can deeply modify species relative abundance, leading to long-lasting changes in community composition. Our study strongly suggests that future research should try to quantify changes in species interactions and shifts in ecological networks resulting from multiple anthropogenic stressors using numerical-based approaches. Furthermore, our findings about species richness and ecosystem functioning relationships force to think about how protection of biodiversity and ecosystem function may require adopting different protection measures. The invaluable objective of biodiversity conservation might not be enough to assure adequate levels of ecosystem functioning.

## Acknowledgements

This study has been funded by the H2020-MSCA-ITN ECORISK2050 project (Grant Agreement nr. 813124) and the CICLIC-ECOREST project (RTI2018-097158-A-C32). A. Rico is supported by a postdoctoral grant provided by the Spanish Ministry of Science, Innovation and University (IJCI – 2017-33465). We thank María José Villena Alvarez (Laboratorios Tecnológicos de Levante) for her contribution to the phytoplankton identification and counting.

## Data accessibility statement

the authors state that, should the manuscript be accepted, the data supporting the results will be archived in an appropriate public repository (Dryad, Figshare or Hal) and the data DOI will be included at the end of the article.

