## Supplementary Info for "Changes in community composition determine recovery trajectories from multiple agricultural stressors in freshwater ecosystems"

#### Pesticide fate

Neither diuron nor chlorpyrifos residues were found in the controls at any time during the experiment, nor traces of diuron were found in mesocosms treated with chlorpyrifos only and vice versa. The DT50 calculated for chlorpyrifos ranged between 3.4 d and 1.6 d. The calculated DT50 for diuron ranged between 45 d and 34 d within the different treatments (Tables S1 and S2). Reasonably, these differences are justified by the different environmental conditions. Faster pesticides' dissipation was noticed in mesocosms treated with nutrients addition. The half-life for chlorpyrifos and diuron in the eutrophic mesocosms were 1.6 and 34 days, respectively, determining a substantial reduction of exposure time to pesticides.

**Table S1.** Relationship between measured and nominal concentrations measured 2h after treatment application, calculated dissipation rate coefficient (k) and half-life value (DT50) for Diuron. Values are expressed as averages with standard deviations between brackets. N: nutrients; CPF: chlorpyrifos; DIU: diuron.

| Treatment | Measured concentration as percentage of nominal (%) | k ($d^{-1}$) | DT50 (d) |
|---|---|---|---|
| CPFxDIU | 115.9 (2.0) | 0.015 | 44.9 |
| CPFxDIUxN | 106.2 (1.9) | 0.019 | 36.7 |
| DIU | 121.2 (4.6) | 0.017 | 40.9 |
| DIUxN | 114.1 (1.7) | 0.020 | 34.2 |

**Table S2.** Relationship between measured and nominal concentrations measured 2h after treatment application, calculated dissipation rate coefficient (k) and half-life value (DT50) for Chlorpyrifos. Values are expressed as averages with standard deviations between brackets. N: nutrients; CPF: chlorpyrifos; DIU: diuron.

| Treatment | Measured concentration as percentage of nominal (%) | k ($d^{-1}$) | DT50 (d) |
|---|---|---|---|
| CPFxDIU | 92.3 (3.9) | 0.21 | 3.4 |
| CPFxDIUxN | 93.1 (6.4) | 0.32 | 2.2 |
| CPF | 90.7 (3.6) | 0.31 | 2.3 |
| CPFxN | 89.1 (6.4) | 0.43 | 1.6 |

### Metabolism

**Table S3.** Mean value and standard deviation for Temperature: Temp. (°C); Dissolved oxygen: DO (mg/L); Oxygen saturation: DO (%), pH (-); Conductivity: EC (µs/cm) and Total Dissolved Solids: TDS (ppm), Total nitrogen: TN (mg/L), and Total phosphate: TP (µg/L) measured during the experimental period in the different treatments. N: nutrients; I: chlorpyrifos; H: diuron.

| Day -5 relative to the treatment | | | | | | | | | | | | | | | | |
|---|---|---|---|---|---|---|---|---|---|---|---|---|---|---|---|---|
| | Temp. (°C) | | DO (mg/L) | | DO (% sat.) | | pH (-) | | EC (µs/cm) | | TDS (ppm) | | TN (mg/L) | | TP (µg/L) | |
| Treat. | mean | ±sd | mean | ±sd | mean | ±sd | mean | ±sd | mean | ±sd | mean | ±sd | mean | ±sd | mean | ±s.d. |
| Control | 17.04 | 0.07 | 14.68 | 0.33 | 163.4 | 5.5 | 9.69 | 0.12 | 189.7 | 9.14 | 98.33 | 10.12 | 0.182 | 0.06 | 45.3 | 23.9 |
| N | 16.96 | 0.1 | 17.31 | 1.22 | 189.5 | 11.9 | 10.03 | 0.15 | 238.33 | 11.95 | 119.33 | 11.68 | 0.31 | 0.17 | 141 | 127 |
| H | 17.06 | 0.05 | 15.64 | 0.19 | 173.3 | 1.7 | 10.38 | 0.08 | 211.67 | 16.65 | 105.67 | 7.77 | 0.158 | 0.04 | 59.3 | 23 |
| HxN | 16.99 | 0.16 | 17.6 | 0.96 | 192.8 | 6.9 | 10.29 | 0.03 | 243.33 | 13.65 | 243.33 | 6 | 0.272 | 0.06 | 129 | 32.6 |
| I | 17.01 | 0.08 | 16.24 | 0.67 | 176.7 | 10 | 10.3 | 0.33 | 232.33 | 30.24 | 116 | 15.39 | 0.283 | 0.15 | 140 | 87.2 |
| IxN | 17.02 | 0.06 | 18.65 | 0.8 | 203.2 | 5.7 | 10.55 | 0.06 | 249.67 | 23.12 | 125 | 11.27 | 0.248 | 0.08 | 150 | 70.6 |
| IxH | 17.02 | 0.08 | 14.36 | 1.12 | 159.8 | 14.4 | 9.99 | 0.24 | 206 | 24.27 | 102.67 | 12.42 | 0.23 | 0.04 | 44.2 | 27.5 |
| IxHxN | 17.04 | 0.23 | 18.64 | 0.94 | 204.7 | 8.4 | 9.19 | 1.01 | 227.67 | 28.36 | 114 | 13.89 | 0.281 | 0.1 | 123 | 66.9 |

| Day 7 relative to the treatment | | | | | | | | | | | | | | | | |
|---|---|---|---|---|---|---|---|---|---|---|---|---|---|---|---|---|
| | Temp. (°C) | | DO (mg/L) | | DO (% sat.) | | pH (-) | | EC (µs/cm) | | TDS (ppm) | | TN (mg/L) | | TP (µg/L) | |
| Treat. | mean | ±sd | mean | ±sd | mean | ±sd | mean | ±sd | mean | ±sd | mean | ±sd | mean | ±sd | mean | ±sd |
| Control | 21.95 | 0.09 | 12.74 | 0.31 | 152.13 | 2.58 | 9.83 | 0.17 | 202.67 | 16.01 | 101 | 8 | 0.201 | 0.0243 | 46.2 | 3.65 |
| N | 21.77 | 0.27 | 14.96 | 0.93 | 182.3 | 10.51 | 10.07 | 0.1 | 261.67 | 10.51 | 164 | 55.97 | 1.16 | 0.174 | 268 | 116 |
| H | 22.21 | 0.42 | 7.12 | 0.28 | 88.77 | 4.12 | 9.02 | 0.04 | 193.67 | 72.6 | 113.67 | 7.77 | 0.194 | 0.0105 | 50.2 | 27.9 |
| HxN | 21.65 | 0.36 | 9.19 | 0.51 | 112.23 | 6.26 | 9.26 | 0.09 | 254.33 | 11.85 | 254.33 | 0.58 | 1.26 | 0.179 | 388 | 166 |
| I | 21.87 | 0.11 | 11.94 | 0.8 | 146.67 | 11.03 | 10.13 | 0.29 | 242 | 38.51 | 154.67 | 74.57 | 0.3 | 0.0793 | 88.8 | 41 |
| IxN | 21.74 | 0.12 | 15.81 | 0.9 | 190.27 | 9.65 | 10.57 | 0.03 | 266.33 | 23.86 | 128.33 | 20.31 | 0.965 | 0.11 | 278 | 92.4 |
| IxH | 22 | 0.12 | 7.45 | 0.76 | 90.97 | 10.26 | 8.64 | 0.37 | 229.33 | 16.17 | 114.67 | 8.08 | 0.221 | 0.01 | 37.3 | 17.6 |
| IxHxN | 21.91 | 0.06 | 9.28 | 0.56 | 113.1 | 6.6 | 9.17 | 0.07 | 243 | 23.26 | 121.67 | 11.68 | 1.1 | 0.0516 | 238 | 82.4 |

**Table S3.** Continued

| Day 15 relative to the treatment | | | | | | | | | | | | | | | | |
|---|---|---|---|---|---|---|---|---|---|---|---|---|---|---|---|---|
| | Temp. (°C) | | DO (mg/L) | | DO (% sat.) | | pH (-) | | EC (µs/cm) | | TDS (ppm) | | TN (mg/L) | | TP (µg/L) | |
| Treat. | mean | ±sd | mean | ±sd | mean | ±sd | mean | ±sd | mean | ±sd | mean | ±sd | mean | ±sd | mean | ±sd |
| Control | 18.65 | 0.19 | 12.39 | 0.8 | 142.77 | 9.03 | 10.07 | 0.08 | 210 | 13.23 | 105 | 6.24 | 0.154 | 0.0333 | 51.2 | 24 |
| N | 18.56 | 0.14 | 13.06 | 0.78 | 150.23 | 7.05 | 10.13 | 0.02 | 256 | 7.05 | 128 | 7.55 | 1.21 | 0.0547 | 312 | 125 |
| H | 18.5 | 0.15 | 9.27 | 0.76 | 106.53 | 8.2 | 8.8 | 0.16 | 260.33 | 14.36 | 128.33 | 9.87 | 0.203 | 0.0625 | 126 | 66.5 |
| HxN | 18.54 | 0.27 | 8.11 | 1.5 | 93.17 | 16.87 | 8.53 | 0.58 | 298.33 | 25.48 | 298.33 | 12.5 | 1.55 | 0.225 | 409 | 98 |
| I | 18.41 | 0.26 | 12.22 | 0.42 | 139.27 | 6.14 | 10.12 | 0.31 | 245.67 | 30.14 | 122.67 | 15.63 | 0.166 | 0.0505 | 69.4 | 19.5 |
| IxN | 18.53 | 0.13 | 12.85 | 0.71 | 147.13 | 8.45 | 10.31 | 0.26 | 261.33 | 17.67 | 130.67 | 9.29 | 1.17 | 0.161 | 315 | 163 |
| IxH | 18.58 | 0.14 | 8.82 | 0.89 | 100.83 | 11.81 | 8.87 | 0.14 | 253.67 | 19.22 | 126.67 | 9.07 | 0.178 | 0.0138 | 46.4 | 13.6 |
| IxHxN | 18.56 | 0.08 | 10.78 | 0.91 | 124.63 | 10.11 | 9.39 | 0.25 | 258 | 28.62 | 196 | 46.68 | 1.35 | 0.0443 | 273 | 86.2 |
| Day 30 relative to the treatment | | | | | | | | | | | | | | | | |
| | Temp. (°C) | | DO (mg/L) | | DO (% sat.) | | pH (-) | | EC (µs/cm) | | TDS (ppm) | | TN (mg/L) | | TP (µg/L) | |
| Treat. | mean | ±sd | mean | ±sd | mean | ±sd | mean | ±sd | mean | ±sd | mean | ±sd | mean | ±sd | mean | ±sd |
| Control | 22.5 | 0.15 | 11.88 | 0.77 | 149.83 | 10.08 | 9.74 | 0.12 | 150.67 | 13.43 | 75.67 | 6.43 | 0.205 | 0.0272 | 34.3 | 21.6 |
| N | 22.42 | 0.13 | 11.96 | 0.42 | 148.33 | 5.3 | 9.65 | 0.15 | 188.33 | 5.3 | 94.33 | 4.04 | 0.692 | 0.0289 | 271 | 106 |
| H | 22.4 | 0.23 | 9.66 | 0.37 | 119.67 | 4.39 | 9.17 | 0.04 | 182.67 | 21.03 | 91.33 | 10.02 | 0.202 | 0.0248 | 118 | 80.3 |
| HxN | 22.31 | 0.07 | 8.08 | 1.74 | 100.47 | 21.9 | 8.34 | 0.64 | 237.33 | 17.62 | 237.33 | 9.02 | 1.47 | 0.414 | 344 | 111 |
| I | 22.32 | 0.2 | 10.2 | 0.98 | 126.63 | 11.91 | 9.72 | 0.32 | 172 | 17 | 86 | 9 | 0.224 | 0.064 | 44.9 | 16.2 |
| IxN | 22.36 | 0.14 | 10.19 | 2.61 | 125.7 | 31.79 | 9.67 | 0.36 | 182.33 | 16.62 | 91.33 | 8.08 | 0.684 | 0.0185 | 292 | 160 |
| IxH | 23.13 | 0.2 | 7.76 | 0.84 | 98.23 | 11.11 | 8.57 | 0.42 | 299.66 | 45.52 | 150 | 21.51 | 0.22 | 0.0226 | 27 | 11.6 |
| IxHxN | 22.48 | 0.16 | 9.91 | 1.7 | 123 | 20.7 | 9.11 | 0.26 | 195.67 | 14.57 | 98 | 7.55 | 0.983 | 0.191 | 264 | 83.3 |

**Table S3.** Continued

| Day 50 relative to the treatment | | | | | | | | | | | | | | | | |
|---|---|---|---|---|---|---|---|---|---|---|---|---|---|---|---|---|
| | Temp. (°C) | | DO (mg/L) | | DO (% sat.) | | pH (-) | | EC (µs/cm) | | TDS (ppm) | | TN (mg/L) | | TP (µg/L) | |
| Treat. | mean | ±sd | mean | ±sd | mean | ±sd | mean | ±sd | mean | ±sd | mean | ±sd | mean | ±sd | mean | ±sd |
| Control | 23.09 | 0.19 | 9.01 | 0.587 | 144.73 | 58.49 | 9.39 | 0.114 | 252.33 | 20.55 | 126 | 10.44 | 0.201 | 0.0165 | 100 | 31.6 |
| N | 22.93 | 0.27 | 97.567 | 0.546 | 121.97 | 5.636 | 9.6 | 0.238 | 289.67 | 5.64 | 145 | 9.54 | 0.507 | 0.0696 | 273 | 55.9 |
| H | 23.22 | 0.05 | 77.467 | 0.705 | 97.633 | 8.425 | 8.4 | 0.463 | 334.67 | 18.58 | 167.33 | 9.29 | 0.21 | 0.0177 | 228 | 60.2 |
| HxN | 23.19 | 0.11 | 12.39 | 0.227 | 154.63 | 3.156 | 9.68 | 0.101 | 339.67 | 15.01 | 339.67 | 31.79 | 0.624 | 0.11 | 331 | 11.9 |
| I | 22.83 | 0.2 | 97.567 | 0.481 | 122.3 | 5.567 | 9.4 | 0.22 | 286.67 | 25.38 | 145.67 | 10.69 | 0.231 | 0.0242 | 120 | 48.3 |
| IxN | 22.87 | 0.31 | 93.933 | 0.981 | 117.8 | 11.96 | 9.6 | 0.175 | 293.67 | 31.18 | 146.67 | 15.31 | 0.47 | 0.0975 | 283 | 51.1 |
| IxH | 23.13 | 0.19 | 77.567 | 0.841 | 98.233 | 11.11 | 8.6 | 0.425 | 299.67 | 43.52 | 150 | 21.52 | 0.185 | 0.0336 | 56.9 | 21.5 |
| IxHxN | 23.46 | 0.1 | 10.57 | 1.805 | 133.4 | 22.21 | 8.9 | 0.665 | 327 | 40.95 | 163.33 | 20.65 | 0.654 | 0.185 | 302 | 69.9 |

**Table S4.** Summary of p-values resulting from the three-way ANOVA test on functional parameters affected by treatments. ↑ indicates a significant increase of the values respect to controls, ↓ indicates a significant decrease of the values respect to controls. '***' p<0.001, '**' p<0.01, '*'p< 0.05. N: nutrients; I: chlorpyrifos; H: diuron.

| | | **N** | **I** | **H** | **IxN** | **HxN** | **IxH** | **IxHxN** |
|---|---|---|---|---|---|---|---|---|
| **pH** | D-5 | 0.850 | 0.370 | 0.630 | 0.400 | 0.770 | 0.350 | 0.540 |
| | D7 | < **0.001** ***↑ | 0.290 | < **0.001** ***↓ | 0.130 | 0.730 | < **0.001** ***↓ | 0.760 |
| | D15 | 0.280 | **0.020** *↑ | < **0.001** ***↓ | 0.060 | 0.980 | 0.140 | 0.160 |
| | D30 | 0.140 | 0.330 | < **0.001** ***↓ | 0.050. | 0.350 | 0.320 | 0.070 |
| | D50 | **0.002** ** ↑ | 0.340 | < **0.001** ***↓ | 0.130 | 0.050 | 0.390 | 0.099 |
| **Chl-a** | | | | | | | | |
| **(µg/L)** | D-5 | **0.005** **↑ | 0.480 | 0.700 | 0.450 | 0.580 | 0.860 | 0.630 |
| | D7 | **0.014** * ↑ | 0.210 | **0.011** * ↑ | 0.730 | < **0.001** ***↓ | 0.880 | 0.160 |
| | D15 | **0.006** **↑ | 0.820 | 0.480 | 0.450 | 0.470 | 0.540 | 0.710 |
| | D30 | 0.079 | 0.550 | 0.720 | 0.980 | 0.740 | 0.460 | 0.740 |
| | D50 | 0.990 | 0.440 | 0.970 | 0.290 | 0.490 | 0.270 | 0.880 |
| **Oxygen %** | D-5 | < **0.001** ***↑ | 0.100 | 0.890 | 0.095 | 0.430 | 0.067 | 0.100 |
| | D7 | < **0.001*****↑ | 0.680 | < **0.001** ***↓ | 0.380 | 0.051 | 0.970 | 0.290 |
| | D15 | 0.140 | 0.270 | < **0.001*****↓ | **0.039** * ↑ | 0.770 | 0.07 . | **0.042** * ↑ |
| | D30 | 0.320 | 0.630 | **0.011** *↓ | 0.630 | 0.400 | **0.011** *↓ | 0.660 |
| | D50 | **0.036** * ↑ | 0.260 | 0.430 | 0.770 | **0.002** **↑ | 0.990 | 0.370 |
| **EC (µs/cm)** | D-5 | **0.004** **↑ | 0.386 | 0.618 | 0.251 | 0.710 | 0.064 | 0.510 |
| | D7 | **0.012***↑ | 0.235 | 0.358 | 0.159 | 0.873 | 0.727 | 0.826 |
| | D15 | **0.009****↑ | 0.866 | **0.013***↑ | 0.086 | 0.589 | **0.023***↑ | 0.925 |
| | D30 | < **.001*****↑ | 0.154 | **0.002****↑ | **0.036***↑ | 0.236 | **0.015***↑ | 0.907 |
| | D50 | 0.120 | 0.844 | **0.001****↑ | 0.866 | 0.800 | 0.084 | 0.276 |

**Phytoplankton**

**Table S5**. Calculated p-values from the three-way ANOVA test. ↑ indicates a significant increase, and ↓ indicates a significant decrease respect to controls. '***' p<0.001, '**' p<0.01, '*'p< 0.05. N: nutrients; CPF: chlorpyrifos; DIU: diuron

| | N | I | H | IxN | HxN | IxH | IxHxN |
|---|---|---|---|---|---|---|---|
| Abundance | | | | | | | |
| D-5 | **0.001**↑** | 0.798 | 0.806 | 0.489 | 0.987 | 0.664 | 0.200 |
| D7 | 0.706 | **0.004**↑** | 0.881 | 0.960 | 0.787 | 0.111 | 0.063 |
| D15 | 0.638 | **0.038*↑** | 0.560 | 0.822 | 0.412 | 0.261 | 0.808 |
| D50 | 0.202 | 0.685 | 0.929 | 0.457 | 0.790 | 0.109 | 0.306 |
| Richness | | | | | | | |
| D-5 | 0.195 | 0.658 | 0.752 | 0.127 | 0.063 | 0.415 | 0.348 |
| D7 | 1.000 | 0.536 | **0.004**↓** | 0.901 | 0.459 | 0.709 | 0.536 |
| D15 | **0.003**↓** | 0.879 | 0.241 | 0.213 | 0.144 | 0.639 | 0.198 |
| D50 | **0.007 **↓** | 0.675 | 0.636 | 0.412 | 0.920 | **0.016 * ↑** | 0.887 |
| Sannon index | | | | | | | |
| D-32 | 0.081 | 0.054 | 0.598 | 0.848 | 0.891 | 0.658 | 0.053 |
| D15 | 0.254 | **0.028*↓** | 0.214 | 0.881 | 0.246 | 0.764 | **0.021*↓** |
| D30 | 0.943 | 0.336 | 0.310 | 0.821 | 0.584 | 0.933 | 0.869 |
| D50 | 0.777 | 0.908 | 0.869 | 0.573 | 0.556 | 0.109 | 0.135 |
| Berger-Parker index | | | | | | | |
| D-32 | 0.097 | 0.090 | 0.709 | 0.882 | 0.960 | 0.699 | 0.071 |
| D15 | 0.076 | **0.004**↑** | 0.392 | 0.780 | 0.111 | 0.981 | **0.001**↑** |
| D30 | 0.776 | 0.952 | 0.418 | 0.869 | 0.870 | 0.684 | 0.500 |
| D50 | 0.989 | 0.733 | 0.939 | 0.638 | 0.616 | 0.549 | 0.229 |

**Table S6**. Summary of p-values for the single and combined effects of different pesticides and nutrients addition on the different taxa populations assessed by factorial the three-way ANOVA. Significant ($p \leq 0.05$) effects and of the treatment are shown in bold. Arrows indicate a treatment-related increase (↑) or decrease (↓) respect to controls. N: nutrients; I: chlorpyrifos; H: diuron.

| | | | | | | | |
|---|---|---|---|---|---|---|---|
| | | | **Day -5** | | | | |
| | **N** | **I** | **H** | **IxN** | **HxN** | **IxH** | **IxHxN** |
| Chlorophyta | **0.004****↑ | 0.642 | 0.667 | 0.639 | 0.766 | 0.778 | 0.083 |
| Desmids | 0.257 | 0.988 | 0.246 | 0.110 | 0.363 | 0.211 | 0.779 |
| Dinophyta | 0.280 | 0.270 | 0.599 | 0.926 | 0.477 | 0.997 | 0.219 |
| Cryptophyta | 0.349 | **0.030 ***↓ | 0.187 | 0.739 | 0.227 | 0.125 | 0.731 |
| Diatoms | 0.823 | 0.819 | 0.743 | 0.464 | 0.379 | 0.944 | 0.466 |
| Cyanobacteria | 0.089 | 0.389 | 0.832 | 0.590 | 0.107 | 0.165 | 0.632 |
| | | | **Day 7** | | | | |
| | **N** | **I** | **H** | **IxN** | **HxN** | **IxH** | **IxHxN** |
| Chlorophyta | 0.273 | **0.002****↑ | 0.508 | 0.124 | 0.675 | 0.574 | **0.007****↑ |
| Desmids | 0.740 | 0.636 | 0.118 | 0.313 | 0.137 | 0.785 | 0.468 |
| Dinophyta | **0.014 ***↓ | 0.070 | 0.632 | 0.973 | 0.190 | 0.935 | 0.973 |
| Cryptophyta | 0.597 | 0.932 | **0.011***↓ | 0.553 | 0.835 | 0.835 | 0.553 |
| Diatoms | 0.516 | 0.729 | **<0.001 *****↓ | 0.485 | 0.485 | 0.883 | 0.661 |
| Cyanobacteria | 0.829 | 0.720 | 0.796 | 0.212 | 0.850 | 0.525 | 0.922 |

**Table S6**. Continued

| Day 15 | | | | | | | |
|---|---|---|---|---|---|---|---|
| | **N** | **I** | **H** | **IxN** | **HxN** | **IxH** | **IxHxN** |
| Chlorophyta | 0.646 | **0.037***↑ | 0.607 | 0.269 | 0.798 | 0.427 | 0.823 |
| Desmids | 0.202 | 0.542 | 0.076 | 0.725 | **0.009****↑ | 0.184 | 0.06504 . |
| Dinophyta | **0.015 ***↓ | 0.417 | 0.725 | 0.146 | 0.417 | 0.723 | 0.146 |
| Cryptophyta | 0.732 | 0.592 | **0.015***↓ | 0.463 | 0.330 | 0.621 | 0.244 |
| Diatoms | 0.638 | 0.310 | 0.054 | 0.673 | 0.539 | 0.931 | 0.970 |
| Cyanobacteria | 0.307 | 0.835 | 0.694 | 0.963 | 0.383 | 0.396 | 0.445 |
| **Day 50** | | | | | | | |
| | **N** | **I** | **H** | **IxN** | **HxN** | **IxH** | **IxHxN** |
| Chlorophyta | 0.457 | 0.690 | 0.913 | 0.181 | 0.514 | 0.772 | 0.437 |
| Desmids | **0.009****↓ | 0.866 | 0.051 | **0.006****↑ | 0.367 | 0.989 | 0.061 |
| Dinophyta | 0.332 | 0.332 | 0.332 | 0.332 | 0.332 | 0.332 | 0.332 |
| Cryptophyta | 0.168 | 0.681 | 0.458 | 0.251 | 0.543 | 0.271 | 0.525 |
| Diatoms | 0.256 | 0.127 | 0.707 | 0.519 | 0.971 | **0.044***↓ | 0.140 |
| Cyanobacteria | 0.198 | 0.261 | **0.044***↓ | 0.084 | 0.867 | 0.251 | 0.750 |

**Figure S1**. Percentage of relative taxa abundance of phytoplankton during the experimental period. C: control; N: nutrients; CPF: chlorpyrifos; DIU: diuron.

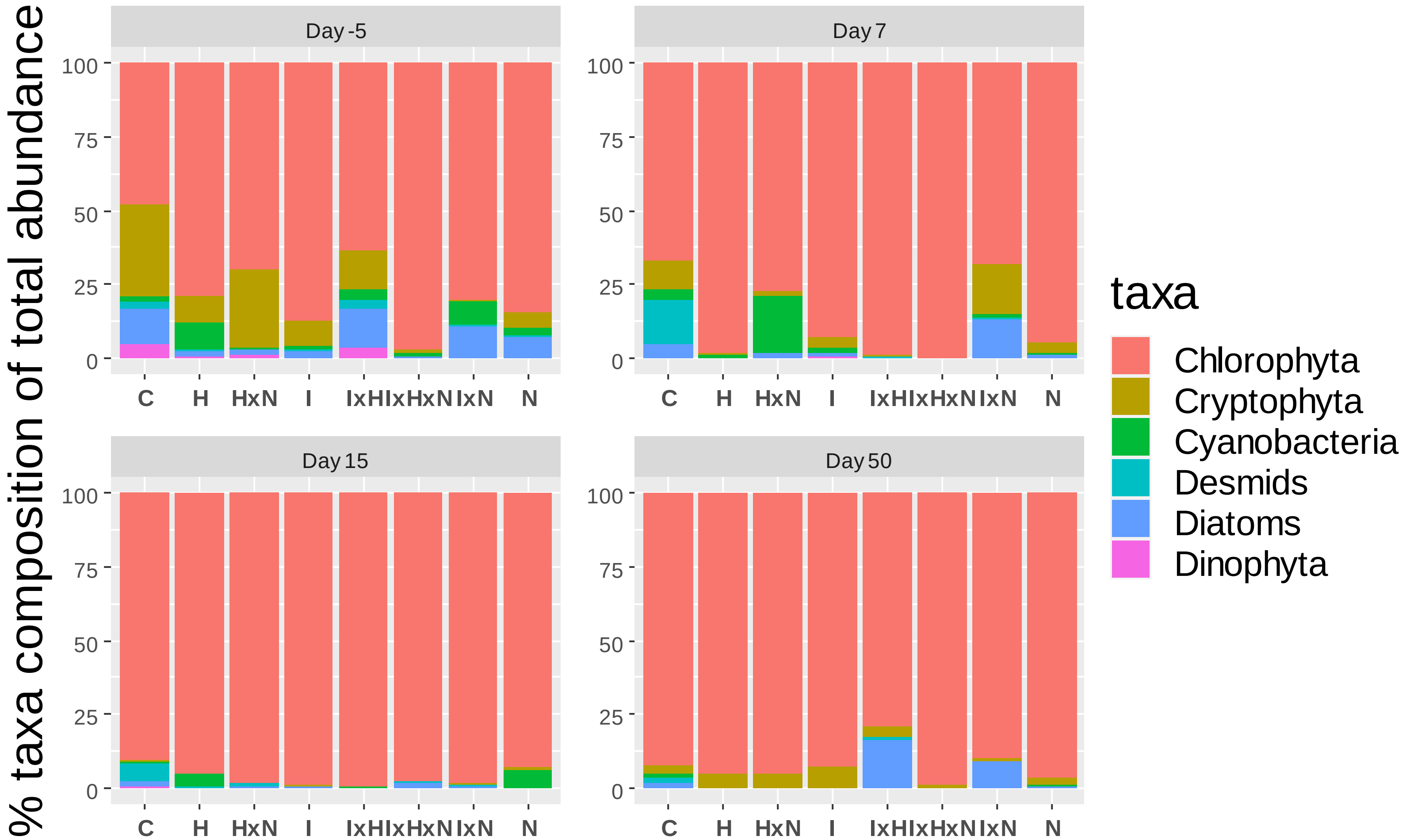

**Figure S2.** Abundance of the Chlorophyta and Diatoms across the experimental period. Data shown are ln-transformed. C: control; N: nutrients; I: chlorpyrifos; H: diuron.

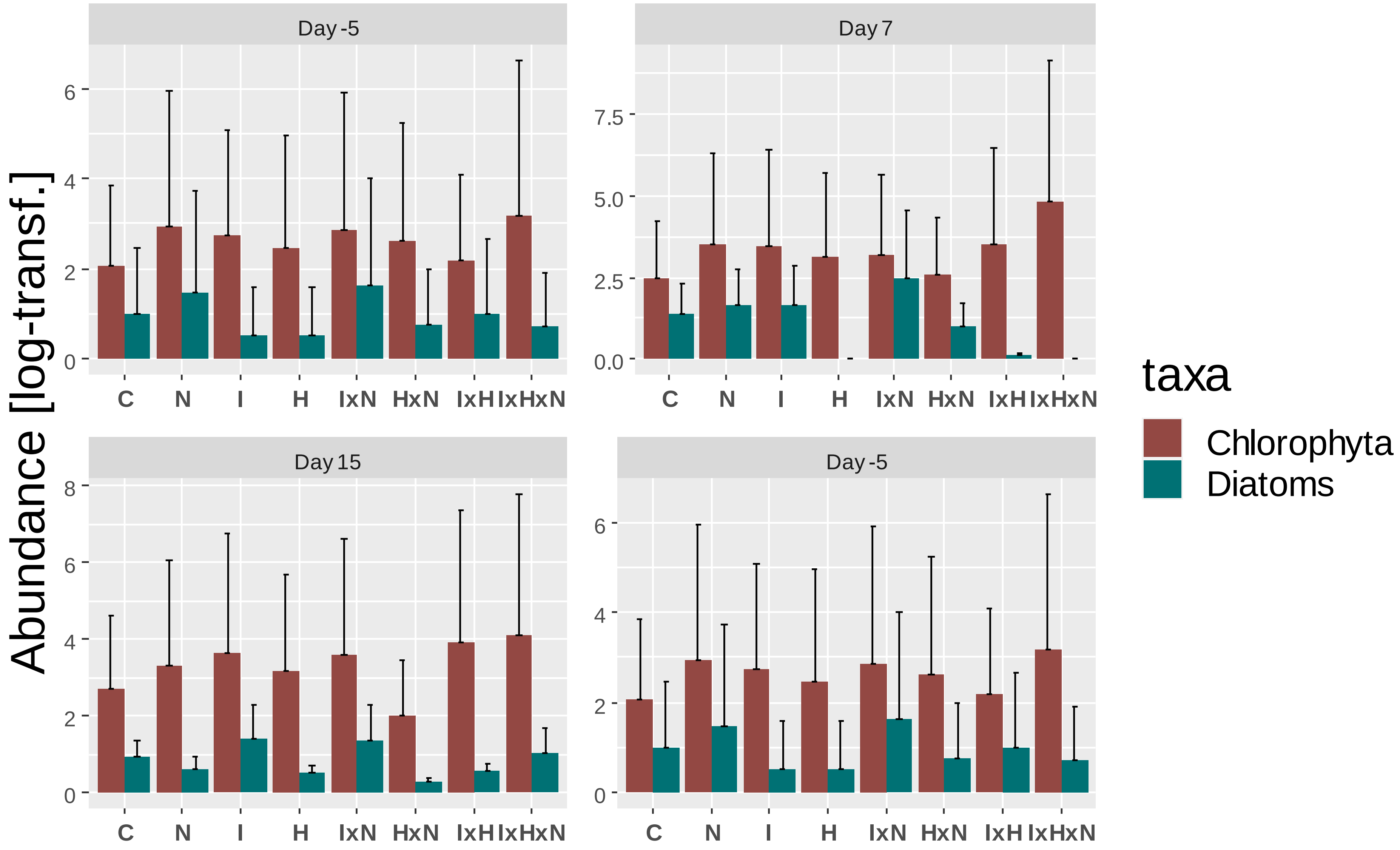

### **Zooplankton**

**Table S7**. Calculated p-values from the three-way ANOVA test on diversity indices. ↑ indicates a significant increase, ↓ indicates a significant decrease respect to controls. '***' p<0.001, '**' p<0.01, '*'p< 0.05. N: nutrients; I: chlorpyrifos; H: diuron.

| | **N** | **I** | **H** | **IxN** | **HxN** | **IxH** | **IxHxN** |
|---|---|---|---|---|---|---|---|
| Abundance | | | | | | | |
| D-5 | 0.563 | 0.267 | 0.441 | 0.161 | 0.224 | 0.589 | 0.719 |
| D7 | 0.550 | 0.208 | 0.623 | **< .001*****↓ | 0.106 | 0.056 | 0.231 |
| D15 | 0.103 | **< .001*****↓ | 0.194 | **0.014***↓ | 0.079 | 0.093 | 0.274 |
| D30 | **0.049***↑ | 0.219 | 0.071 | 0.287 | 0.408 | 0.505 | 0.875 |
| D50 | 0.431 | 0.236 | 0.640 | 0.184 | 0.549 | 0.214 | 0.443 |
| Richness | | | | | | | |
| D-5 | 0.661 | 0.137 | 0.826 | 0.826 | 0.512 | 0.661 | 0.661 |
| D7 | 0.402 | 0.529 | 0.115 | 0.097 | 0.109 | 0.192 | 0.186 |
| D15 | **0.016 *** ↓ | **0.009****↓ | 0.567 | 0.308 | 0.098 | 0.425 | 0.730 |
| D30 | **0.036***↓ | **< .001*****↓ | **0.011 *** ↓ | 0.170 | 1.000 | **0.021***↓ | 0.400 |
| D50 | **0.018 *** ↓ | **< .001*****↓ | 0.077 | 0.461 | 0.710 | **0.0081 **** ↓ | 0.461 |
| Shannon index | | | | | | | |
| D-5 | 0.560 | 0.932 | 0.173 | 0.230 | 0.501 | 0.085 | 0.751 |
| D7 | 0.671 | 0.413 | 0.556 | **<.001 *****↑ | **0.042 *** ↓ | 0.401 | 0.726 |
| D15 | **<.001 *****↓ | **0.010 *** ↓ | **0.041 *** ↑ | **0.034 *** ↑ | **0.029 ***↓ | 0.989 | 0.518 |
| D30 | **0.008****↓ | **0.0026 ****↓ | 0.297 | 0.724 | 0.919 | 0.341 | 0.922 |
| D50 | 0.083 | **0.0018 **** ↓ | 0.715 | **0.011 *** ↓ | 0.270 | 0.183 | 0.667 |
| Berger-Parker index | | | | | | | |
| D-5 | 0.625 | 0.846 | 0.261 | 0.660 | 0.522 | 0.123 | 0.930 |
| D7 | 0.274 | 0.534 | 0.554 | **< .001 *****↓ | 0.087 | 0.091 | 0.500 |
| D15 | **0.003 ****↑ | | 0.062 | 0.151 | 0.052 | 0.795 | 0.396 |
| D30 | **0.02 *** ↑ | **0.006 ****↑ | 0.387 | 0.401 | 0.294 | 0.892 | 0.699 |
| D50 | 0.189 | **0.004 ****↑ | 0.380 | **0.006 ****↑ | 0.352 | 0.903 | 0.466 |

**Table S8.** Summary of p-values for the single and combined effects of different pesticides and nutrients addition on the different zooplankton populations assessed by the three-way ANOVA. Significant ($p \leq 0.05$) effects of the treatment are shown in bold. Arrows indicate a treatment-related increase (↑) or decrease (↓) respect to controls. N: nutrients; I: chlorpyrifos; H: diuron.

| | | | **Day -5** | | | | |
|---|---|---|---|---|---|---|---|
| | **N** | **I** | **H** | **IxN** | **HxN** | **IxH** | **IxHxN** |
| **Cladocera** | | | | | | | |
| Daphnia sp | 0.350 | 0.480 | 0.480 | 0.190 | 0.064 | 0.210 | 0.710 |
| Ceriodaphnia | 0.630 | 0.190 | 0.088 | 0.650 | 0.220 | 0.360 | 0.360 |
| Alonella | 0.690 | 0.590 | 0.120 | 0.160 | 0.790 | 0.120 | 0.360 |
| Simocephalus | 0.940 | 0.480 | 0.460 | 0.440 | 0.260 | 0.230 | 0.100 |
| **Copepoda** | | | | | | | |
| Cyclopodia | **0.003****↓ | 0.690 | 0.790 | 0.120 | 0.110 | 0.400 | 0.430 |
| Nauplii | 0.990 | 0.350 | 0.510 | 0.450 | 0.500 | 0.370 | 0.330 |
| **Rotifera** | | | | | | | |
| Testudinella | 0.400 | 0.160 | 0.140 | 0.640 | 0.450 | **0.001****↑ | 0.580 |
| Polyartha | 0.300 | 0.990 | 0.450 | 0.190 | 0.600 | 0.540 | 0.570 |
| | | | **Day 7** | | | | |
| | **N** | **I** | **H** | **IxN** | **HxN** | **IxH** | **IxHxN** |
| **Cladocera** | | | | | | | |
| Daphnia sp | 0.059 | **0.032 ***↓ | 0.840 | 0.530 | **0.034 ***↓ | 0.790 | 0.940 |
| Ceriodaphnia | 0.770 | **0.003****↓ | 0.990 | 0.660 | 0.370 | 0.330 | 0.310 |
| Alonella | 0.630 | **<0.001 *****↓ | 0.680 | 0.330 | 0.930 | 0.490 | 0.220 |
| Simocephalus | 0.061 | 0.740 | 0.590 | 0.052 | **0.036 ***↓ | 0.450 | 0.290 |
| **Copepoda** | | | | | | | |
| Cyclopodia | 0.570 | 0.170 | 0.550 | 0.260 | 0.720 | 0.300 | 0.460 |
| Nauplii | 0.091 | **0.001****↓ | 0.220 | 0.680 | **0.022 *** ↓ | 0.960 | 0.740 |
| **Rotifera** | | | | | | | |
| Testudinella | 0.061 | 0.620 | 0.160 | 0.100 | 0.083 | 0.096 | 0.170 |
| Polyartha | **<0.001 *****↓ | **<0.001 *****↑ | **0.003**** ↓ | 0.960 | **0.001****↓ | **0.005**** ↓ | **0.011*** ↓ |

**Table S8**. Continued

| | N | I | H | IxN | HxN | IxH | IxHxN |
|---|---|---|---|---|---|---|---|
| | | | **Day 15** | | | | |
| | **N** | **I** | **H** | **IxN** | **HxN** | **IxH** | **IxHxN** |
| **Cladocera** | | | | | | | |
| Daphnia sp | **0.011** *↑ | 0.270 | 0.840 | 0.730 | 0.300 | 0.330 | 0.530 |
| Ceriodaphnia | 0.560 | **<0.001*****↓ | 0.940 | 0.800 | 0.450 | 0.790 | 0.990 |
| Alonella | 0.560 | **<0.001*****↓ | 0.680 | 0.680 | 0.400 | 0.740 | 0.740 |
| Simocephalus | 0.450 | 0.230 | 0.900 | 0.320 | 0.890 | 0.072 | 0.084 |
| **Copepoda** | | | | | | | |
| Cyclopodia | 0.250 | **0.001****↑ | 0.480 | 0.260 | **0.026*** ↑ | 0.120 | 0.130 |
| Nauplii | **0.001**** ↑ | **<0.001** *****↓ | **0.012*** ↑ | **0.012***↓ | **<0.001** *****↓ | **0.032** * ↑ | **0.023*** ↓ |
| **Rotifera** | | | | | | | |
| Testudinella | **0.001****↓ | 0.360 | 0.440 | 0.300 | 0.100 | 0.570 | 0.220 |
| Polyartha | **0.012** *↓ | 0.490 | 0.170 | 0.340 | 0.540 | 0.270 | 0.500 |
| | | | **Day 30** | | | | |
| | **N** | **I** | **H** | **IxN** | **HxN** | **IxH** | **IxHxN** |
| **Cladocera** | | | | | | | |
| Daphnia sp | 0.091 | 0.140 | 0.270 | 0.220 | 0.069 | 0.320 | 0.610 |
| Ceriodaphnia | 0.710 | 0.055 | 0.730 | 0.800 | 0.580 | 0.420 | 0.580 |
| Alonella | 0.690 | **0.002****↓ | 0.110 | 0.110 | 0.690 | 0.370 | 0.370 |
| Simocephalus | 0.160 | 0.700 | **0.006**↓ | 0.930 | 0.550 | 0.770 | 0.870 |
| **Copepoda** | | | | | | | |
| Cyclopodia | 0.370 | 0.330 | 0.860 | **0.042** *↓ | 0.540 | 0.084 | 0.980 |
| Nauplii | **0.018** *↑ | 0.080 | 0.150 | 0.570 | 0.260 | 0.500 | 0.770 |
| **Rotifera** | | | | | | | |
| Testudinella | 0.690 | **0.002****↓ | 0.110 | 0.110 | 0.690 | 0.370 | 0.370 |
| Polyartha | 0.360 | 0.330 | 0.860 | **0.042** *↓ | 0.540 | 0.084 | 0.980 |

**Table S8**. Continued

| | Day 50 | | | | | | |
|---|---|---|---|---|---|---|---|
| | **N** | **I** | **H** | **IxN** | **HxN** | **IxH** | **IxHxN** |
| **Cladocera** | | | | | | | |
| Daphnia sp | 0.290 | 0.260 | **0.045** *↓ | **0.016** *↓ | 0.830 | **0.042** *↓ | 0.430 |
| Ceriodaphnia | 0.250 | **<0.001** ***↓ | 0.094 | **0.008**** ↓ | 0.052 | 0.320 | 0.180 |
| Alonella | 0.950 | 0.170 | 0.820 | 0.290 | 0.230 | **0.007****↓ | 0.320 |
| Simocephalus | 0.830 | 0.910 | 0.270 | 0.650 | 0.190 | 0.067 | 0.270 |
| **Copepoda** | | | | | | | |
| Cyclopodia | 0.091 | 0.240 | 0.860 | 0.110 | 0.550 | **0.021** *↓ | 0.760 |
| Nauplii | 0.860 | 0.350 | 0.510 | 0.690 | 0.320 | 0.620 | 0.430 |
| **Rotifera** | | | | | | | |
| Testudinella | **0.024** *↓ | **0.046** *↓ | 0.620 | 0.280 | 0.500 | 0.350 | 0.730 |
| Polyartha | 0.180 | 0.970 | 0.180 | 0.980 | 0.980 | 0.180 | 0.980 |

**Figure S3**. Zooplankton abundance on day 30. C: control; N: nutrients; I: chlorpyrifos; H: diuron.

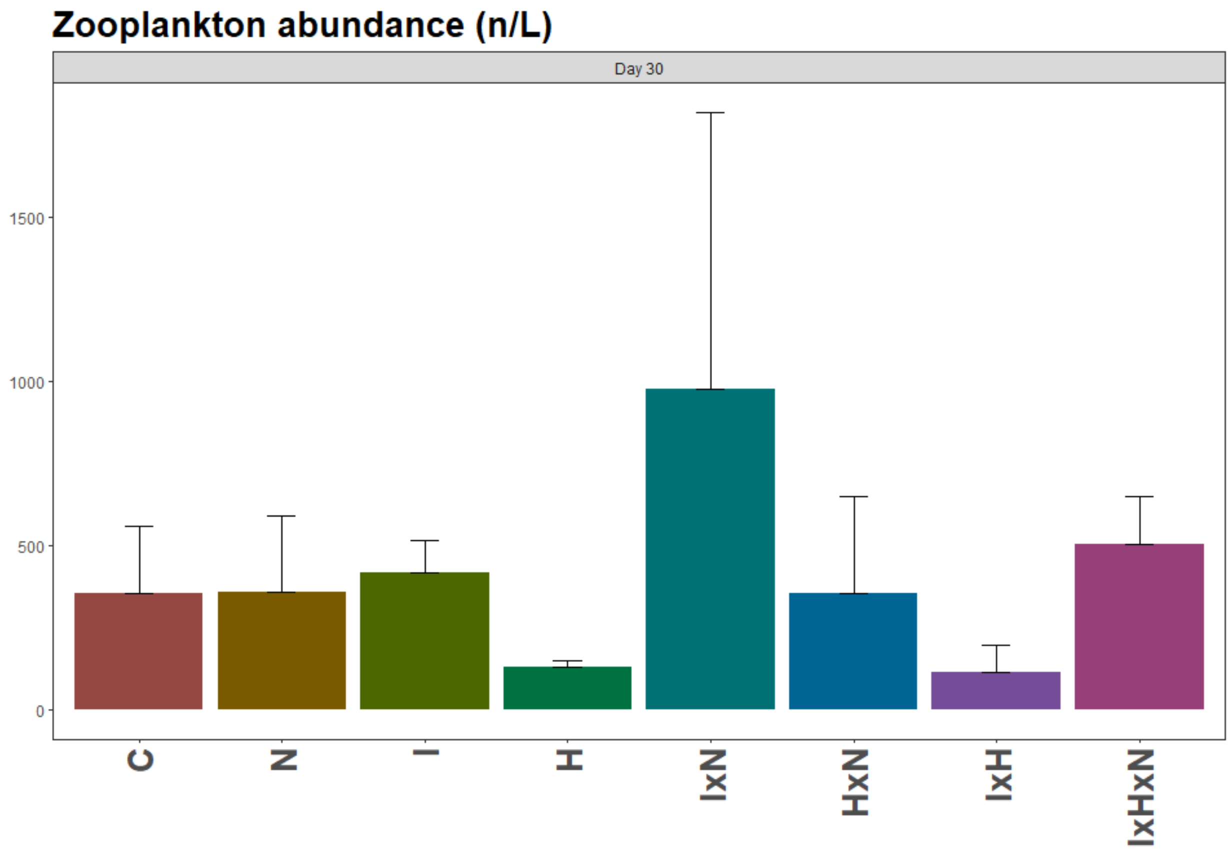

**Figure S4**. Percentage of relative taxa abundance of zooplankton across the experimental period. C: control; N: nutrients; I: chlorpyrifos; H: diuron.

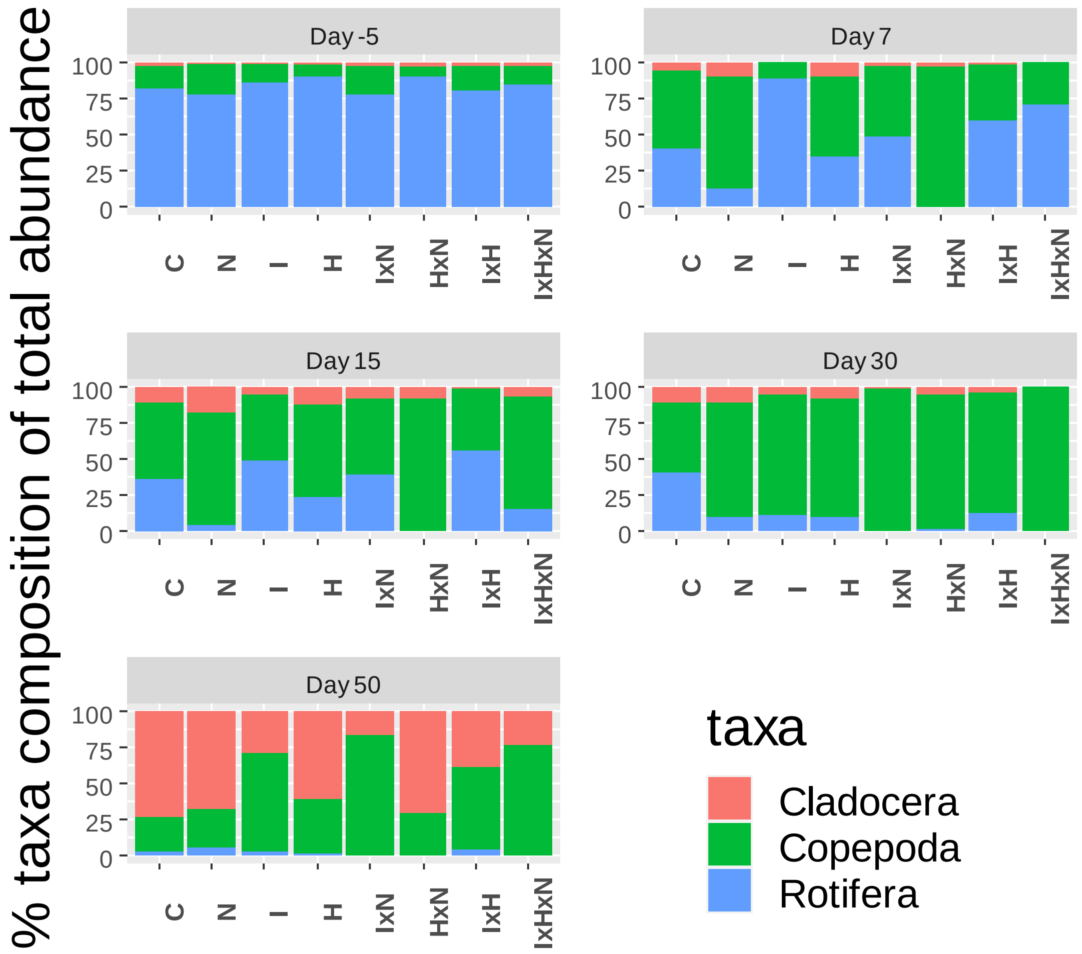

**Figure S5.** Abundance of Copepoda and Cladocera across the experimental period. C: control; N: nutrients; I: chlorpyrifos; H: diuron.

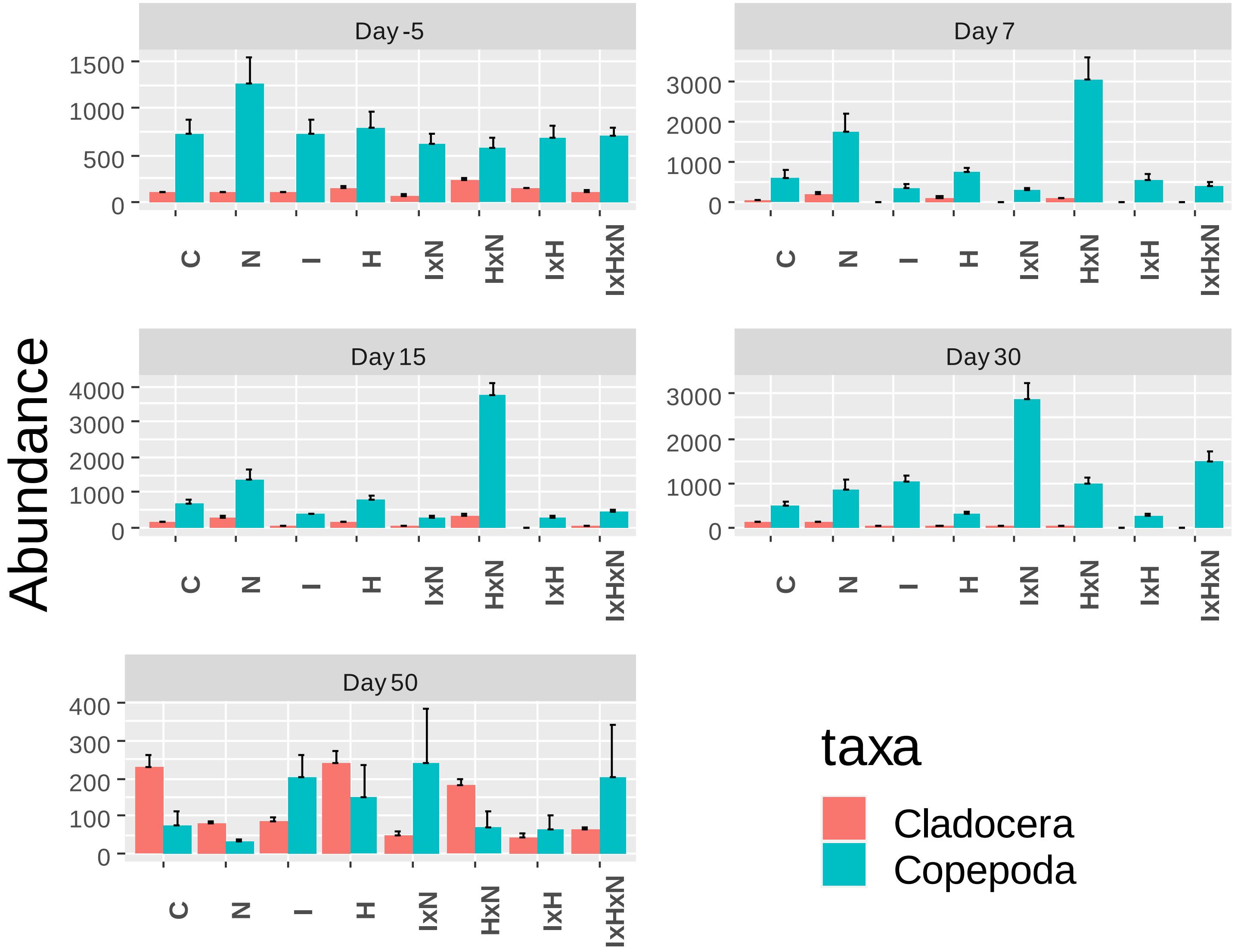

**Macroinvertebrates**

**Table S9.** Calculated p-values from the three-way ANOVA test on diversity indices. ↑ indicates a significant increase of the values, ↓ indicates a significant decrease of the values respect to controls. '***' p<0.001, '**' p<0.01, '*'p< 0.05. N: nutrients; I: chlorpyrifos; H: diuron

| | **N** | **I** | **H** | **IxN** | **HxN** | **IxH** | **IxHxN** |
|---|---|---|---|---|---|---|---|
| Abundance | | | | | | | |
| D-32 | 0.910 | 0.870 | 0.143 | 0.075 | 0.760 | 0.990 | 0.110 |
| D15 | **0.0054 **↑** | **0.013*↓** | 0.480 | 0.390 | 0.460 | 0.730 | 0.760 |
| D30 | **< .001***↑** | 0.890 | 0.790 | 0.360 | **0.027 *↑** | 0.150 | 0.110 |
| D50 | **0.034 *↑** | 0.350 | 0.200 | 0.600 | 0.490 | 0.220 | 0.590 |
| richness | | | | | | | |
| D-32 | 0.057 | 0.906 | 0.412 | 0.137 | 0.204 | 0.09 | 0.412 |
| D15 | 0.176 | **0.012*↓** | 0.305 | 0.176 | 0.728 | 0.728 | 0.728 |
| D30 | 0.472 | 0.203 | 0.664 | **023*↓** | 0.885 | 0.317 | **0.042*↑** |
| D50 | **0.013*↓** | 0.61 | **0.020 *↑** | 0.45 | 0.80 | 0.61 | 0.80 |
| Shannon index | | | | | | | |
| D-32 | 0.446 | 0.216 | 0.868 | 0.255 | 1 | 0.107 | 0.901 |
| D15 | 0.898 | 0.669 | 0.151 | 0.91 | 0.898 | 0.584 | 0.874 |
| D30 | 0.779 | 0.274 | 0.359 | 0.2 | 0.659 | 0.556 | 0.985 |
| D50 | **0.016 *↓** | 0.93 | 0.35 | 0.07 | 0.89 | 0.48 | 0.44 |
| Berger-Parker index | | | | | | | |
| D-32 | 0.93 | 0.89 | 0.73 | 0.34 | 0.42 | 0.25 | 0.90 |
| D15 | 0.24 | 0.93 | **0.047 *↑** | 0.89 | 0.78 | 0.64 | 0.94 |
| D30 | 0.5694 | 0.3751 | 0.1473 | .2125 | 0.9328 | 0.5019 | 0.3556 |
| D50 | **0.024*↑** | 0.89 | 0.34 | 0.09 | 0.51 | 0.21 | 0.71 |

**Table S10.** Summary of p-values for the single and combined effects of different pesticides and nutrients addition on the different taxa populations assessed by factorial ANOVA. Significant ($p \leq 0.05$) effects of the treatment are shown in bold. Arrows indicate a treatment-related increase (↑) or decrease (↓) respect to controls. N: nutrients; I: chlorpyrifos; H: diuron.

| | | | | | | | | |
|---|---|---|---|---|---|---|---|---|
| **Day -32** | | | | | | | | |
| **Insecta** | | **N** | **I** | **H** | **IxN** | **HxN** | **IxH** | **IxHxN** |
| Diptera | Chironomini | 0.913 | 0.728 | 0.953 | 0.982 | 0.029 | 0.849 | 0.635 |
| | Orthocladiinae | 0.145 | 0.642 | 0.362 | 0.27 | 0.891 | 0.519 | 0.362 |
| Ephemeropetera | Cloeon sp. | 0.366 | 0.181 | 0.386 | 0.575 | 0.443 | 0.144 | 0.647 |
| | Caenidae | 0.132 | 0.475 | 0.406 | 0.344 | 1000 | 0.719 | 0.24 |
| Odonata | Zygoptera | 0.722 | 0.93 | 0.954 | 0.833 | 0.77 | 0.633 | 0.286 |
| Trichoptera | Leptoceridae | 0.967 | 0.208 | 0.208 | 0.901 | 0.433 | 0.433 | 0.102 |
| **Mollusca** | | | | | | | | |
| Mesogastropoda | Hydrobiidae | **0.068↑** | 0.225 | **0.035↑** | **0.015↑** | **0.048↑** | 0.102 | 0.314 |
| Basommatophora | Physela acuta | 0.722 | 0.93 | 0.954 | 0.833 | 0.77 | 0.633 | 0.286 |
| **Anellida** | | | | | | | | |
| Haplotaxida | Lumbriculidae | 0.927 | 0.227 | 0.442 | 0.911 | 0.385 | 0.495 | 0.82 |
| **Tricladida** | | | | | | | | |
| Dugesiidae | Dugesia sp. | 0.658 | 0.38 | 0.068 | 0.463 | 0.463 | 0.768 | 0.556 |
| **Day 15** | | | | | | | | |
| **Insecta** | | **N** | **I** | **H** | **IxN** | **HxN** | **IxH** | **IxHxN** |
| Diptera | Chironomini | **<0.001↑** | **<0.001↓** | 0.897 | 0.397 | 0.263 | 0.898 | 0.347 |
| | Orthocladiinae | 0.761 | 0.881 | **0.007↓** | **0.027** | 0.635 | 0.635 | 0.635 |
| Ephemeropetera | Cloeon sp. | 0.054 | **<0.001↓** | 0.947 | 0.859 | 0.95 | **0.086↓** | 0.856 |
| | Caenidae | **0.093↑** | 1000 | 1 | 0.384 | 0.384 | 0.384 | 1000 |
| Odonata | Zygoptera | 0.398 | 0.729 | 0.369 | 0.611 | 0.493 | 0.358 | 0.548 |
| Trichoptera | Leptoceridae | 0.823 | **0.058↓** | 0.192 | 0.505 | 0.273 | 0.655 | 0.655 |
| **Mollusca** | | | | | | | | |
| Mesogastropoda | Hydrobiidae | 0.498 | 0.331 | 0.374 | 0.642 | **0.027↑** | 0.311 | 0.209 |
| Basommatophora | Physela acuta | 0.2 | 0.459 | 0.455 | 0.782 | 0.278 | 0.469 | 0.848 |
| **Anellida** | | | | | | | | |
| Haplotaxida | Lumbriculidae | 0.888 | 0.914 | 0.369 | 0.189 | 0.466 | 0.123 | 0.203 |
| **Tricladida** | | | | | | | | |
| Dugesiidae | Dugesia sp. | 0.634 | **0.027↑** | **0.007↑** | **0.027** | 0.634 | 0.635 | 0.635 |

**Table S10.** Continued

| Day 30 | | | | | | | | |
|---|---|---|---|---|---|---|---|---|
| **Insecta** | | **N** | **I** | **H** | **IxN** | **HxN** | **IxH** | **IxHxN** |
| Diptera | Chironomini | **0.014**↑ | **0.026**↓ | 0.565 | **0.093**↓ | 0.631 | 0.679 | 0.569 |
| | Orthocladiinae | 0.748 | 0.354 | 0.621 | **0.002**↓ | 0.565 | 0.470 | 0.497 |
| Ephemeropetera | Cloeon sp. | 0.378 | 0.458 | 0.164 | 0.491 | 0.306 | 0.307 | 0.262 |
| | Caenidae | 0.423 | **0.025**↓ | 0.807 | 0.243 | 0.329 | 0.532 | 0.134 |
| Odonata | Zygoptera | 0.181 | 0.64 | 0.405 | 0.587 | 0.332 | 0.843 | 0.766 |
| Trichoptera | Leptoceridae | 0.847 | 0.189 | **0.021**↓ | 0.189 | 0.189 | 0.341 | 0.341 |
| **Mollusca** | | | | | | | | |
| Mesogastropoda | Hydrobiidae | 0.289 | 0.291 | 0.299 | 0.294 | 0.292 | 0.287 | 0.285 |
| Basommatophora | Physela acuta | **0.043**↑ | 0.459 | 0.189 | 0.844 | 0.548 | 0.906 | 0.398 |
| **Anellida** | | | | | | | | |
| Haplotaxida | Lumbriculidae | 0.816 | 0.197 | 0.536 | 0.959 | 0.413 | 0.764 | 0.067 |
| **Tricladida** | | | | | | | | |
| Dugesiidae | Dugesia sp. | **0.039**↑ | **0.047**↑ | 0.300 | **0.004**↑ | **0.039**↑ | **0.047**↑ | 0.300 |
| Day 50 | | | | | | | | |
| **Insecta** | | **N** | **I** | **H** | **IxN** | **HxN** | **IxH** | **IxHxN** |
| Diptera | Chironomini | **0.028**↑ | 0.38 | 0.529 | 0.631 | 0.547 | **0.071**↓ | 0.942 |
| | Orthocladiinae | 0.623 | 0.638 | 0.954 | 0.478 | 0.375 | 0.109 | **0.035**↓ |
| Ephemeropetera | Cloeon sp. | 0.297 | **0.084**↑ | 0.984 | 0.24 | 0.991 | **0.013**↑ | 0.797 |
| | Caenidae | **0.011**↓ | 0.579 | 0.799 | **0.065**↓ | 0.779 | 0.862 | 0.541 |
| Odonata | Zygoptera | 0.482 | 0.81 | 0.298 | 0.757 | 0.43 | 0.941 | 0.313 |
| Trichoptera | Leptoceridae | 0.332 | 0.332 | 0.332 | 0.332 | 0.332 | **0.033**↓ | 0.743 |
| **Mollusca** | | | | | | | | |
| Mesogastropoda | Hydrobiidae | 0.498 | 0.331 | 0.311 | 0.374 | **0.027**↑ | 0.642 | 0.209 |
| Basommatophora | Physela acuta | **0.002** ↑ | 0.934 | 0.773 | 0.32 | 0.271 | 0.888 | 0.735 |
| **Anellida** | | | | | | | | |
| Haplotaxida | Lumbriculidae | 0.504 | 0.16 | 0.364 | 0.3 | 0.325 | 0.66 | 0.347 |
| **Tricladida** | | | | | | | | |
| Dugesiidae | Dugesia sp. | 0.726 | 0.675 | 0.171 | 0.675 | 0.171 | **0.073**↑ | 0.955 |

**Figure S6**. Percentage of relative taxa abundance across the experimental period for macroinvertebrates. C: control; N: nutrients; I: chlorpyrifos; H: diuron.

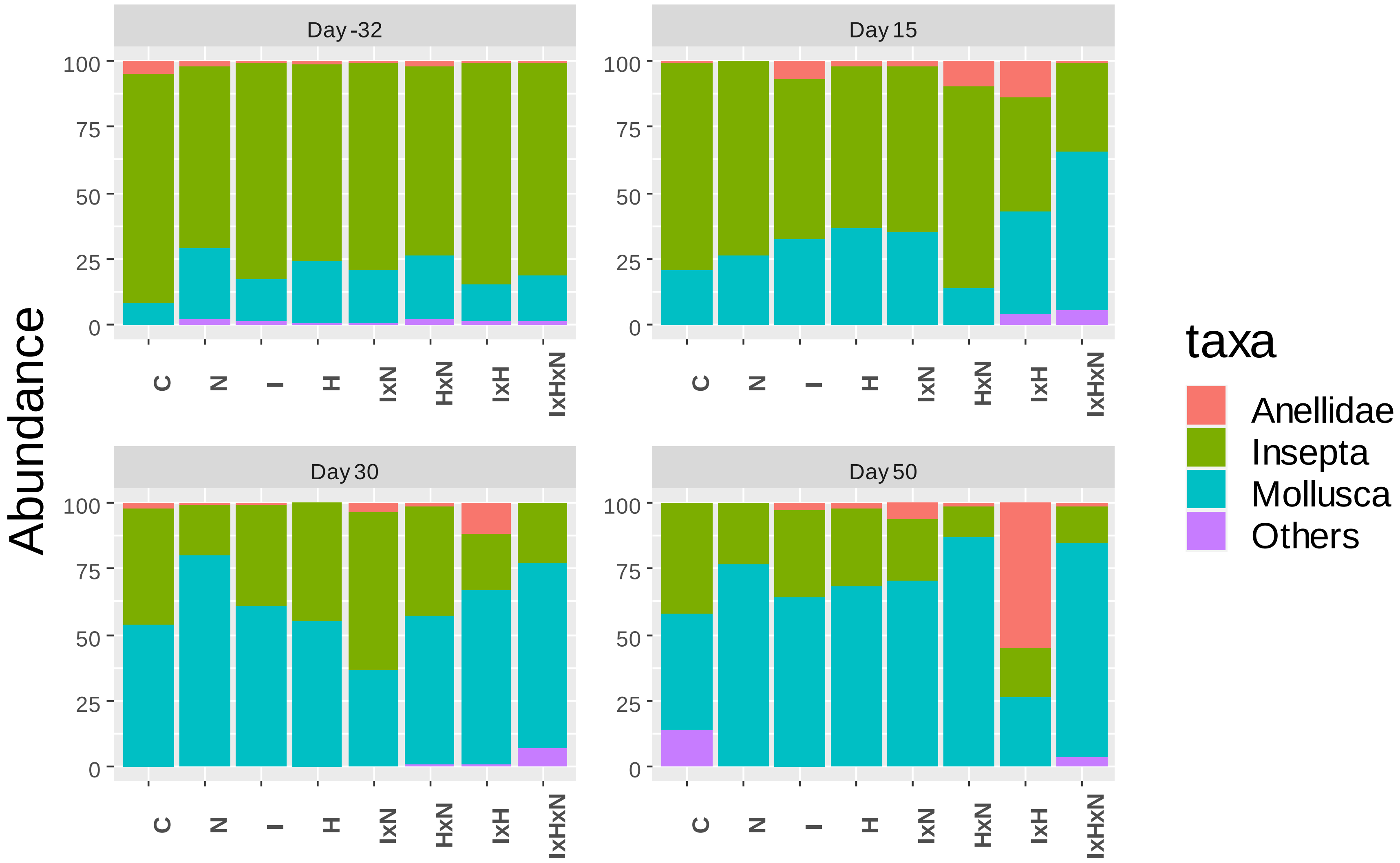

**Figure S7.** Abundance of Chironomini and Ephemeroptera across the experimental period. C: control; N: nutrients; I: chlorpyrifos; H: diuron.

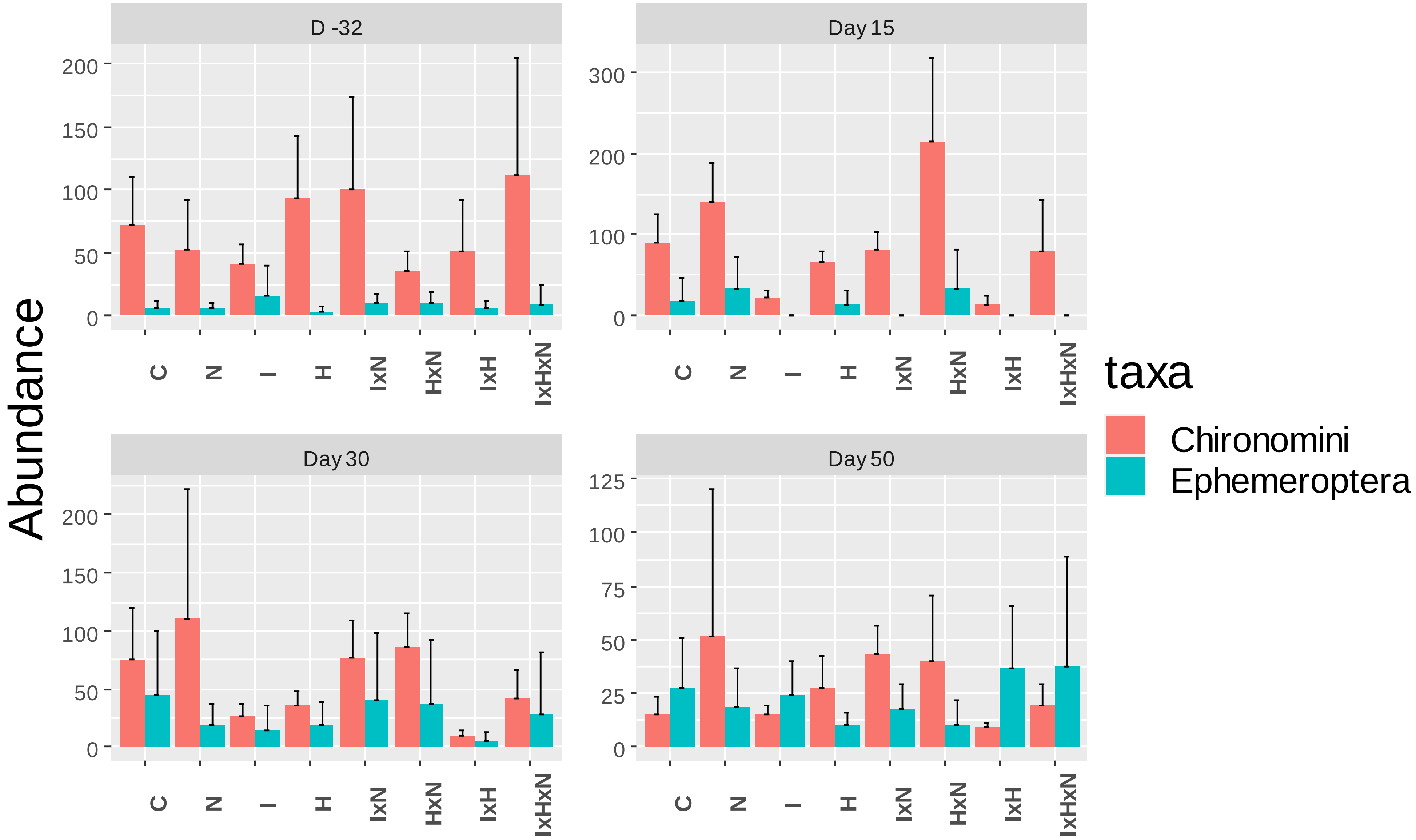

**Figure S8.** . Conceptual map of the combined effects of nutrients, chlorpyrifos and diuron. Arrows indicate a stressor-related increase (↑) or decrease (↓), (-) indicates no effect. Red lines connecting the boxes represent direct effects, and blue lines represent indirect effects.

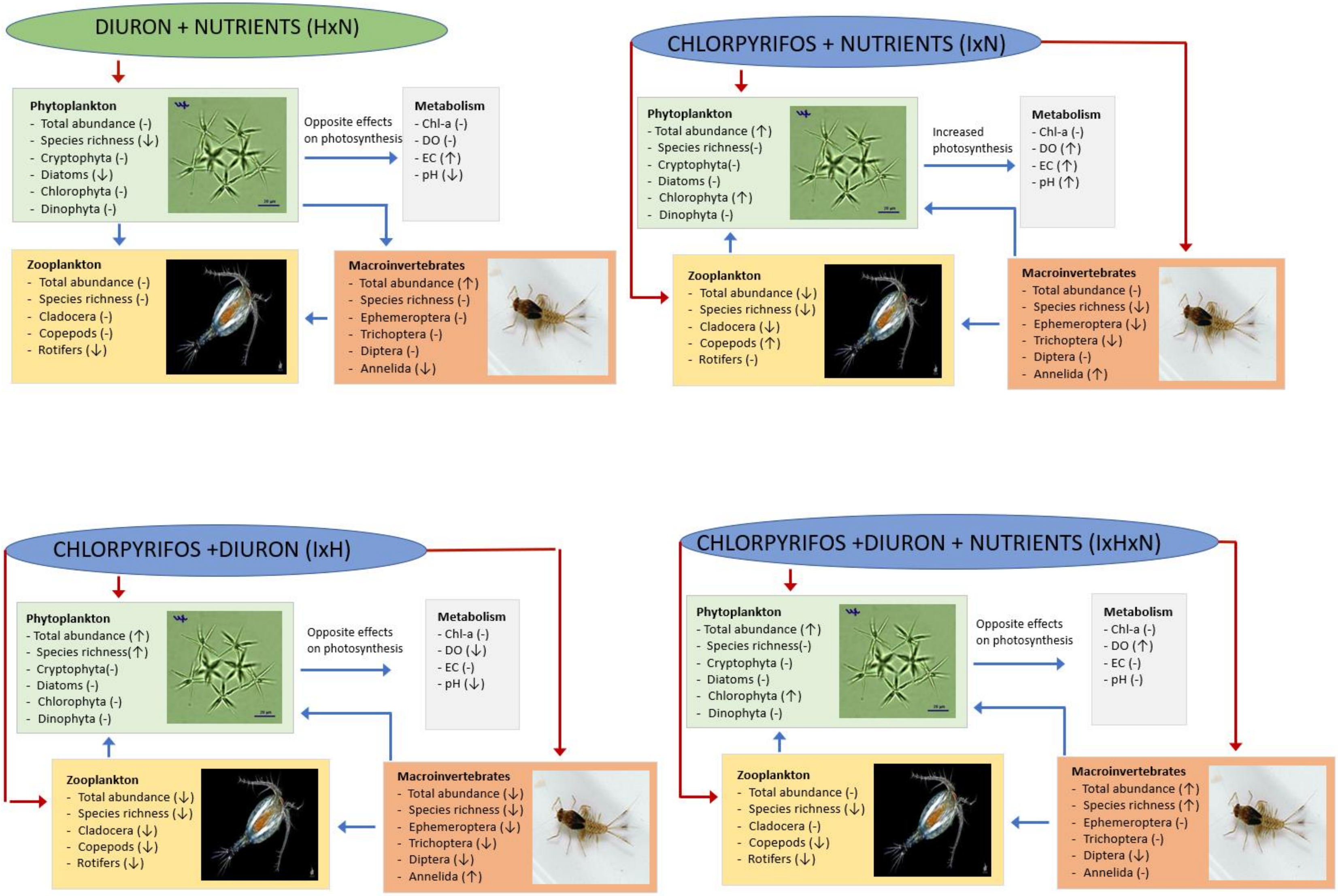
